# The role of concentration on drop formation and breakup of collagen, fibrinogen, and thrombin solutions during inkjet bioprinting

**DOI:** 10.1101/2020.10.06.328187

**Authors:** Hemanth Gudapati, Ibrahim T. Ozbolat

## Abstract

The influence of protein concentration on drop formation and breakup of aqueous solutions of fibrous proteins collagen, fibrinogen, and globular protein thrombin in different concentration regimes is investigated during drop-on-demand (DOD) inkjet bioprinting. The capillary-driven thinning and breakup of dilute (*c/c*\* < 1, where *c* is the concentration and *c*\* is the overlap concentration) collagen, fibrinogen, and thrombin solutions is predominantly resisted by inertial force on the initial onset of necking. The minimum diameter (*D_f_min__*(*t*)) of the necked fluid up to the critical pinch-off time (*t*_*c*_) scales with time as *D_f_min__*(*t*) ∼ (*t_c_* − *t*)^2/3^, a characteristic of potential flows. Although the capillary-driven thinning and breakup of semidilute unentangled collagen (1 ≤ *c/c*\* ≤ 4) and fibrinogen (1 ≤ *c/c*\* ≤ 1.3) solutions is predominantly resisted by inertial force on the initial onset of necking, the breakup of droplets is delayed beyond *t*_*c*_, where the minimum diameter of the necked fluid decreases exponentially with time because of the resistance of elastic force. The resistance of viscous force to the necking of both the dilute and semidilute untangled protein solutions is negligible. Aggregates or subvisible particles (between 1 and 100 μm) constantly disrupt the formation of droplets for the semidilute unentangled protein solutions, even when their inverse Ohnesorge number (*Z*) is within the printability range of 4 ≤ *Z* ≤ 14. Although aggregates are present in the dilute protein solutions, they do not disrupt the formation of droplets.

## Introduction

Drop-on-demand (DOD) inkjet printing accurately delivers small volumes of fluids (between 1 and 100 pL) to fabricate features at a high spatial resolution of ≤ 100 μm for structural and functional material applications.^1,2^ Collagen type I, fibrinogen, and thrombin proteins in aqueous buffer solutions are widely used as precursors of natural biopolymers for tissue engineering applications.^3,4^ Drop-on-demand inkjet three-dimensional (3D) bioprinting of such buffered protein solutions is highly desired for accurately delivering different living cells and biologics (such as growth factors) to recapitulate native tissue physiology during the fabrication of high-throughput disease models and reconstruction of damaged tissues etc. Drop-on-demand inkjet printing relies on acoustic pressure waves resulting from the localized heating of the printed solution (thermal DOD inkjet) or the physical deformation of the inkjet nozzle (piezoelectric DOD inkjet) to form droplets.^2^ During the formation of droplets, a fluid thread or column, commonly referred to as ligament, is projected outwards through the nozzle orifice at first. The outward projection of the ligament depends on inertia or kinetic energy of the acoustic pressure waves in overcoming the viscous friction associated with the shear viscosity of the printed solution and surface tension at the nozzle orifice.^5^ The ligament, once projected outwards, begins to thin and a droplet eventually pinches off under the action of capillary pressure.

Traditionally, surfactants such as Triton-X-100 and viscosity modifiers such as glycerol were used for successfully delivering droplets of various buffered protein solutions using inkjet printing.^6–9^ However, surfactants and viscosity modifiers may not only interfere with the polymerization of collagen and fibrinogen, but are cytotoxic and immunogenic.^10–14^ Hence, inkjet printing of collagen, fibrnogen, and thrombin solutions without the use surfacntants and viscosity modifiers is highly desired for tissue engineering applications. Interestingly, inkjet printing of collagen and thrombin soutions without the use of surfactants and viscosity modifiers has been reported by several research groups. For example, Boland *et al.*^15–17^ have used a commercially available thermal inkjet paper printer to deliver droplets of collagen and thrombin solutions for various biological applications. Similarly, Sanjana *et al.*^18^ and Zarowna-Dabrowska *et al.*^19^ have used piezoelectric inkjet printers for delivering droplets of collagen solutions. Unfortunately, inkjet printing of collagen solutions with a concentration > 1 mg/mL and fibrinogen solutions, which are biologically relevant for tissue engineering applications, has not been reported in the literature. Although inkjet printing of thrombin solution with a concentration of ≤ 50 U/mL has been reported,^16^ it is not known whether those solutions had formed stable droplets or unstable satellite droplets.

The viscoelastic response of collagen and fibrinogen solutions increases with concentration and can delay or completely prevent the breakup of droplets.^20^ Moreover, collagen, fibrinogen, and thrombin molecules adsorb and aggregate at the solution/air interface of protein solutions.^20^ Such an aggregation increases with concentration and the aggregated proteins can desorb from the interface because of mechanical disturbances and cause further aggregation in the bulk.^21,22^ Similarly, the proteins may adsorb and aggregate at the solution/solid interface and desorb owing to mechanical disturbances, causing further aggregation in the bulk.^23^ The presence of particles in the printed fluid generally entraps air in the fluid, which dissipates the acoustic pressure waves and disrupts the formation of droplets during printing.^24^ Similarly, the bulk protein aggregates can disrupt the formation of droplets during printing. Hence, an understanding of the dynamics of droplet formation and breakup of the protein solutions in different concentration regimes without the use of surfactants and viscosity modifiers is important for adapting the inkjet printing of those solutions for tissue engineering applications.

The primary objective of this work is to gain a better understanding of the dynamics of droplet formation and breakup of ligaments during DOD inkjet printing of surfactant-free collagen type I, fibrinogen, and thrombin solutions in dilute and semidilute unentangled concentration regimes. The secondary objective of this work is to identify the process parameters for the formation of stable droplets of those protein solutions. This work through sequential photographic images of evolving ligaments and the corresponding ligament diameter measurements proves that the capillary thinning of ligaments of the investigated dilute and semidilute protein solutions during printing is predominantly resisted by inertial force on the initial onset of necking. However, elastic force delays the breakup of the ligaments of the semidilute protein solutions. In this work, aggregates refer to subvisible particles, including but not limited to supra-monomeric assemblies, which are observed near the nozzle orifice during DOD inkjet printing of protein solutions. Such supra-monomeric assemblies may form because of reversible self-association, reversible clustering, and irreversible clustering of the protein monomers.

## Background

### Drop-on-demand inkjet printing modalities

Drop-on-demand inkjet printing generally forms droplets using two different modalities.^2^ The first modality is thermal inkjet printing. This modality forms droplets relying on the acoustic pressure waves resulting from the rapid expansion and collapse of a vapor bubble, arising because of the localized heating of the printed solution. The second modality is piezoelectric inkjet printing. This modality forms droplets relying on the radial deformation of a piezoelectric transducer (see Figure S1 in the Supporting Information). Piezoelectric inkjet printing was used in this study because, during thermal inkjet printing, the temperatures of the printed solution near the heating element far exceeds the melting temperature (*T*_*m*_, the temperature at which 50% of the proteins denature) of collagen and fibrinogen.^25–27^

### Evolution and breakup of ligaments during piezoelectric DOD inkjet printing

Formation of droplets during piezoelectric inkjet printing begins with the actuation of a piezoelectric transducer. The piezoelectric transducer radially expands at first and contracts afterwards (see Figure S1 in the Supporting Information). The initial radial expansion of the transducer produces a negative acoustic pressure (or dilation) wave which propagates in the directions of both open and closed (nozzle orifice) ends.^28–30^ The negative pressure wave, which propagates to the open end, is reflected back as a positive pressure (or compression) wave at the boundary. At the same time, the negative pressure wave, which propagates to the closed end, is reflected back as a negative pressure wave at the boundary. The two reflected pressure waves superpose over each other at a certain time point, resulting in zero total pressure. However, this does not stop the propagation of the waves. At this instance, when the transducer radially contracts back, the negative pressure wave is canceled and the positive pressure wave is amplified. The amplification of the positive pressure wave projects a ligament of the printed solution with the maximum possible inertia or kinetic energy through the orifice and a droplet breaks off with the maximum possible ejection velocity.^5^ The corresponding dwell time (*t*_*dwell*_) for this high energy droplet formation is given by the relationship:^28,30,31^

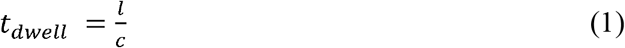

where *l* is the capillary length (which is the length between the open and closed ends) and *C* is the effective speed of sound in the printed fluid.

When the radially expanded piezoelectric transducer does not contract at *l/C*, the negative pressure wave, which is reflected from the closed end, propagates to the open end. At the same time, the positive pressure wave, which is reflected from the open end, propagates to the closed end. The negative pressure wave is reflected back as a positive pressure wave whereas the positive pressure wave is reflected back as a positive pressure wave. The two positive pressure waves superpose over each other at the instant 2(*l/C*), doubling the total pressure. When the piezoelectric transducer radially contracts back at this instant, the superimposed pressure waves are amplified and a ligament is ejected. The ligament has a low kinetic energy when compared to ligament ejected at the instant *l/C*. However, this suppresses the formation of undesired satellite droplets for some fluids.^30^

The evolution and breakup of the ligaments of Newtonian fluids during inkjet printing depends on the delicate balance of inertial, viscous, and capillary forces. Jang *et al*.^32^ studied the influence of viscosity and surface tension on the formation stable droplets of Newtonian fluids with Ohnesorge (*Oh*) number and its inverse value (*Z*) during inkjet printing:

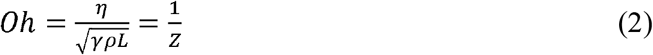

where *η, γ* and *ρ* are the viscosity, surface tension, and density of the fluids, respectively. In addition, *L* is the characteristic length and *U* is the velocity of a droplet. In this study, the nozzle diameter is chosen as the characteristic length. When *Z* < 4 (based on nozzle diameter), the inertia/kinetic energy of the acoustic pressure wave is too small to overcome the viscous friction and the surface tension at the nozzle orifice. As a result, no droplets are ejected. When *Z* > 14, the inertia of the acoustic pressure wave is too large and/or capillary pressure is too dominant during the breakup of the ligament. As a result, multiple satellite droplets are ejected along with a primary droplet. The evolution and breakup of ligaments of viscoelastic fluids during inkjet printing depends on the delicate balance of inertial, viscous, elastic, and capillary forces, in that viscoelasticity delays or prevents the breakup of ligaments.^33,34^

For both Newtonian and Non-Newtonian viscoelastic fluids in a simple extensional flow (such as capillary break up studies by Basaran *et al*.^35^) when *Oh* ≪ 1 (based on nozzle orifice radius), the inertial force dominates and predominantly resists the capillary thinning up to the critical pinch-off time (*t*_*c*_).^36,37^ The critical pinch-off time *t*_*c*_ is the time at which the necked fluid pinches off from rest of the fluid. Under this regime, and the minimum ligament diameter (*D_f_min__*(*t*)) follows the scaling law of: ^34,37,38^

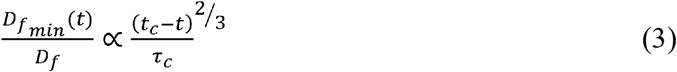

Here, *D*_*f*_ is the initial ligament diameter and *t* is the time, and τ_*c*_ is the inertio-capillary time scale:

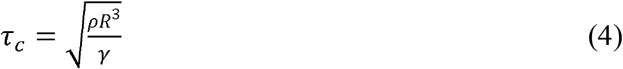

where *R* is the radius of the nozzle orifice. Conversely, when *Oh* ≫ 1, the viscous force dominates and predominantly resists the capillary thinning up to the critical pinch-off time *t*_*c*_ for both Newtonian and Non-Newtonian viscoelastic fluids. Under this regime, and the minimum ligament diameter *D_f_min__*(*t*) follows the scaling law of: ^34,37,38^

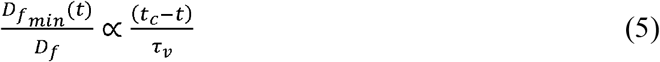

where τ_*v*_ is the visco-capillary time scale:

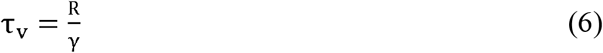

As the critical pinch-off time *t*_*c*_ approaches for Newtonian fluids, both the inertial and viscous forces become important at the pinch-off region, regardless of the viscosity (or *Oh* number) of the fluids.^37^ In addition, a secondary microthread develops at the pinch-off region between the necked fluid and rest of the fluid. The viscous length scale (*l*_*sec*_) of the microthread is given by:

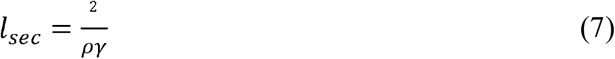

and the viscous time scale (*t*_*sec*_) of the microthread is given by: ^38^

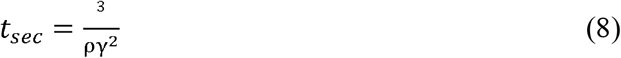

Often, *l*_*sec*_ is in nanometers and *t*_*sec*_ is nanoseconds for fluids such as water because of which the secondary thread is not generally observable.

As the critical pinch-off time *t*_*c*_ approaches for non-Newtonian viscoelastic fluids such as polymer solutions, the tensile elastic stress becomes important. It has been suggested that the elastic stress, arising because of stretching of the polymer chains, resists the capillary thinning and prevents the separation of the necked fluid at the critical pinch-off time *t*_*c*_.^39,40^ A cylindrical microthread develops between the necked fluid and rest of the fluid at the critical pinch-off time *t*_*c*_. The diameter of the microthread decreases exponentially with time and the exponential decrease is inversely proportional to the characteristic relaxation time (*λ*_*ch*_) of the solutions:^40^

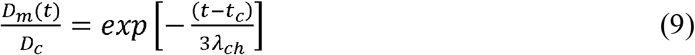

where *D*_*m*_(*t*) is the diameter of the cylindrical microthread and *D*_*c*_ is the diameter of the ligament at the critical pinch-off time *t*_*c*_. The characteristic relaxation time λ_*ch*_ depends on polymer concentration.^34^ At some time point beyond the critical pinch-off time *t*_*c*_, the polymer chains are completely stretched and the elastic resistance to the capillary thinning saturates.^40^ As a result, the ligament diameter decreases linearly with time and eventually a droplet pinches off.

Aqueous sodium alginate solutions during inkjet printing have been shown to follow the scaling laws established for viscoelastic fluids in a simple extensional flow.^34^ The typical diameter of a ligament during the inkjet printing is < 100 μm and the corresponding Bond number (*B*_*0*_ = *ρgL*^*2*^/*γ*, where *g* is acceleration due to gravity) is ≪ 1. Accordingly, the influence of gravity on the ligament pinch-off behavior and spreading of impinging droplets on a substrate is considered negligible.

## Materials and methods

### Protein solutions

Surfactant-free collagen type I, fibrinogen, and thrombin buffered protein solutions were prepared from the very same batches of the protein solutions which were used in our previous study.^20^ The preparation of protein solutions and their viscometry, steady and dynamic shear rheology, overlap concentration (*c*\*, the concentration above which protein monomers begin interpenetrate each other)^41^, intrinsic viscosity ([η]), density, and surface tension measurements were reported in that study. Briefly, collagen type I was extracted from rat tail tendons as freeze-dried sponges according previously published protocols.^42,43^ The extracted collagen sponges were dissolved in 0.02 N acetic acid (cat. no. 695092, Sigma-Aldrich, St. Louis, MO) to prepare a sterile stock collagen solution, which was stored at 4 °C. At the time of printing, the stock collagen solution was neutralized to desired concentrations using sterile 10X Phosphate-Buffered Saline solution (PBS, pH 7.4, cat. no. 46-013-CM, Corning Inc., Corning, NY), sterile 1 N NaOH solution (Sigma-Aldrich) and sterile deionized (DI) water. The neutralized collagen solutions were placed on an ice bath during printing.

The stock fibrinogen solution was prepared by dissolving bovine fibrinogen protein crystals isolated from bovine plasma (cat. no. F8360, Sigma-Aldrich) in Dulbecco’s Phosphate-Buffered Saline solution (DPBS, pH 7.4, cat. no. 20-031-CV, Corning). Similarly, the stock thrombin solution was prepared by dissolving bovine thrombin protein powder isolated from bovine plasma (cat. no. T4648, Sigma-Aldrich) in DPBS. The stock fibrinogen and thrombin solutions were stored at −30 °C and were diluted to desired concentrations with DPBS at the time of printing. The fibrinogen and thrombin solutions were maintained at room temperature during printing. In addition, collagen and fibrinogen solutions were filtered with sterile syringe filters to prepare filtered protein solutions at the time of printing. The collagen solutions were filtered with a 0.45 μm pore size syringe filter (Biomed Scientific) whereas fibrinogen solutions were filtered with a 0.22 μm pore size syringe filter (cat. no. SLGPM33RS, Sigma-Aldrich).

### Piezoelectric DOD inkjet printing experiments

A custom-built inkjet printer (jetlab® 4, MicroFab Technologies, Plano, TX) was used for studying drop formation and breakup of the protein during inkjet printing. The printer consisted of four piezoelectric dispensing devices (120 μm nozzle orifice diameter, cat. no. MJ-ABL-01-120-8MX, MicroFab Technologies) and accordingly the three different proteins were printed with seperate dispensing devices to avoid cross contamination. The inkjet printer also consisted of different controllers for controlling the actuation voltage pulse, movement of the dispensing devices/printhead assembly, and back-pressure (*BP*) of the dispensing devices. A simple unipolar trapezoidal voltage pulse was chosen to actuate the dispensing devices to minimize experimental variables. A positive back-pressure was initially used for supplying protein solutions from the reservoirs to the dispensing devices. Afterwards, a negative back-pressure was used for preventing the dripping of the solutions from the device. A software program provided by the manufacturer was used for operating the controllers. The printer was also equipped with a strobe light illuminator and a horizontally mounted microscopic camera (resolution: 640H x 480V, Sentech STC-MB33USB, Sensor Technologies America, Carrollton, TX) to observe the formation of droplets. The droplet formation images were captured with an open-source screen recording program (CamStudio, CamStudio.org).

The sequential droplet formation images were captured by delaying the illumination of the strobe light at the time steps of 1 μs after the actuation of each voltage pulse. That is, the first image was taken when the strobe light was illuminated 0 μs after the actuation of the first voltage pulse. Whereas, the second image was taken when the strobe light was illuminated 1 μs after the actuation of the second voltage pulse and so on. The first and second (and third and so on) voltage pulses were separated by a time interval of 0.25 – 1 s, and between 25 and 30 images were taken in total. Hence, the representative sequential droplet formation images of a single droplet were essentially stitched from the images of multiple ligaments/droplets. Those images were analyzed with an open-source image analysis software program (ImageJ, National Institutes of Health, Bethesda, MD) to characterize the diameter and velocity (see Figure S1 in the Supporting Information) of the ligaments and droplets. The piezoelectric dispensing devices were regularly cleaned by filling them with regular household bleach (The Clorox Company, Oakland, CA) or Dimethylformamide (DMF, Sigma-Aldrich) and placing them in an ultrasonic cleaning bath for 30 min.

Semidilute unentangled collagen solutions with a concentration (*c*) between 1 and 2 mg/mL had a *Z* value between 4 and 15 and were predicted to form stable droplets with a 120 μm nozzle orifice during inkjet printing.^20,32^ Conversely, dilute collagen solutions with *c* ≤ 0.5 mg/mL, dilute and semidilute unentangled fibrinogen solutions with *c* ≤ 50 mg/mL, and dilute thrombin solutions with *c* ≤ 20 U/mL had a *Z* value between 15 and 100 and were predicted to form multiple satellite droplets with a 120 μm nozzle orifice.^20^ However, the formation of stable droplets of water with a *Z* value of ~ 100 during inkjet printing was successfully demonstrated in multiple studies.^44–46^ Accordingly, a frequency (*f*) between 20 and 200 Hz was used for all the investigated protein solutions. In addition, Milli-Q water was used as a control fluid in this study (see Figures S2-7 in the Supporting Information for more information on inkjet printing of Milli-Q water).

## Results and discussion

### Drop-on-demand inkjet printing of surfactant-free collagen solutions

Droplet formation images of dilute collagen solutions with concentrations of 0.3 mg/mL (*c/c*\* = 0.59) and 0.5 mg/mL (*c/c*\* = 0.99) and semidilute unentangled collagen solutions with concentrations of 1 mg/mL (*c/c*\* = 2), and 2 mg/mL (*c/c*\* = 4) and the corresponding actuation voltage pulse properties are presented in Figure 1. The collagen solutions with concentrations of 0.3 and 0.5 mg/mL form stable droplets, which is in agreement with a previous study by Sanjana *et al*,^18^ whom produced stable droplets with 0.5 mg/mL collagen solutions. Conversely, the collagen solutions with concentrations of 1 and 2 mg/mL form unstable droplets, where it was difficult to identify optimal dwell time because of the constant presence of aggregates. The dwell time (*t*_*dwell*_) for the investigated collagen solutions is 50 μs. This is in agreement with the theoretical dwell time of 2(*l/C*) for a less energetic droplet formation, which suppresses the formation of satellite droplets. Because the solutions are in dilute and semidilute unentangled concentration regimes, the speed of sound in those solutions is comparable to speed of sound in Milli-Q water,^47^ such that 2*l/C* (*l* ~ 30 mm and *C* =1450 m/s in water) ≈ 50 μs.

**Figure 1.**
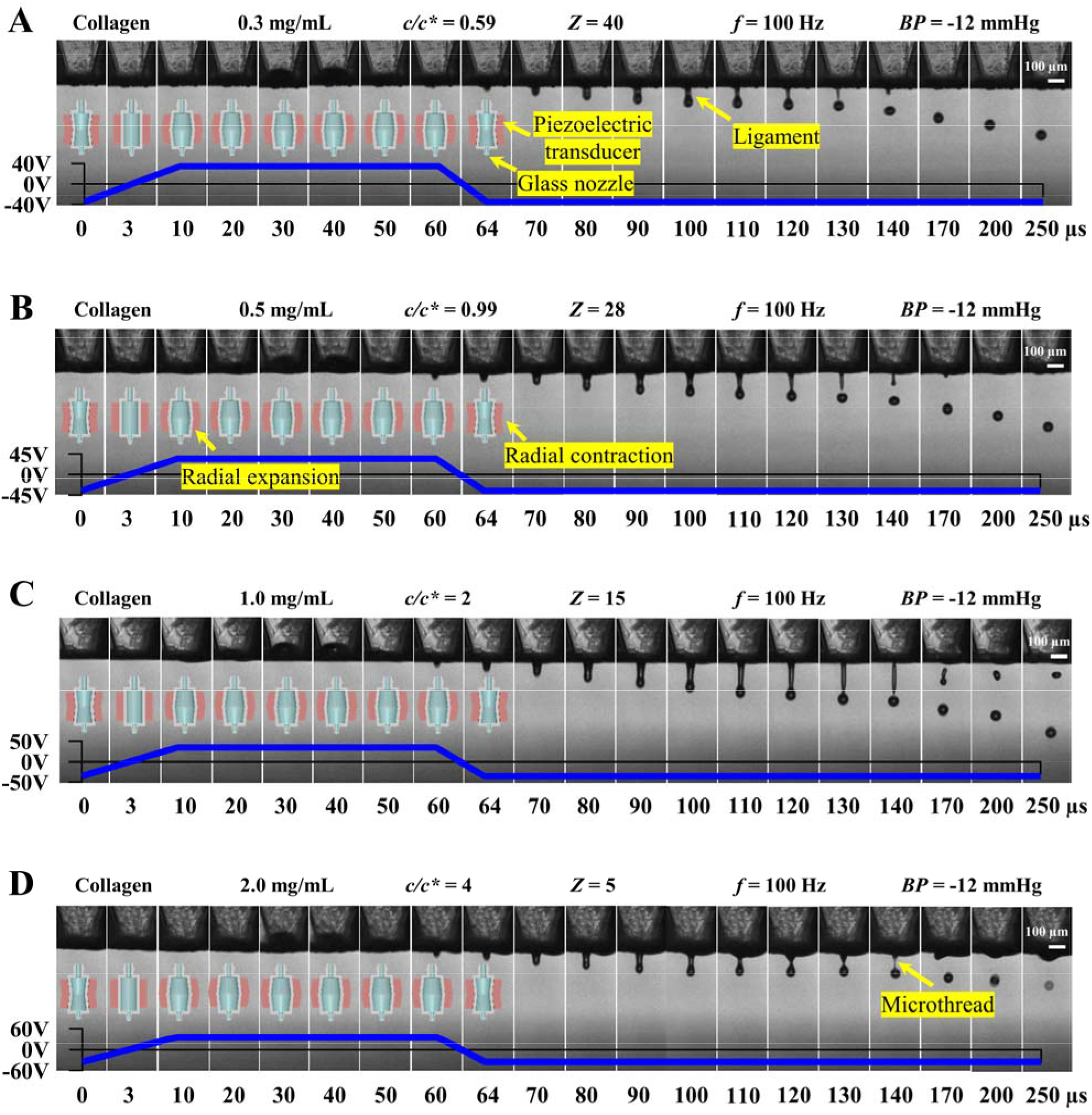
Drop-on-demand inkjet printing of dilute and semidilute unentangled collagen solutions. (A) 0.3 mg/mL, (B) 0.5 mg/mL, (C) 1 mg/mL, and (D) 2 mg/mL collagen solutions. The *Z* values are based on the nozzle orifice diameter. The sequential representative image showing single droplet formation, which were assembled by capturing images of different droplets at different time points.

The ligaments emerge out of the orifice at 60 μs which is similar to Milli-Q water. However, droplets pinch off at 130 – 140 μs, which is 20 – 30 μs longer when compared to those of Milli-Q water ejected with a back pressure of −18 mmHg. The elastic force can delay the capillary thinning of the ligaments of collagen solutions. In addition, the ligament diameters of collagen solutions, which are ejected with a back pressure of −12 mmHg, are 5 – 10 μm larger than ligament diameter of Milli-Q water. In general, the ligament diameter and the corresponding ligament pinch-off time increase with decreasing back pressure (see Figures S3-4 in the Supporting Information). Thus, the ligament pinch-off for collagen solutions takes longer because of their viscoelasticity and larger ligament diameters when compared to Milli-Q water.

The diameter of ligaments as a function of time is presented in Figure 2. The minimum ligament diameter *D_f_min__*(*t*) scales as (*t*_*c*_ – *t*)/τ_*c*_ to the power of 2/3 up to the critical pinch-off time *t*_*c*_ for all protein concentrations. Accordingly, inertial force predominantly resists the capillary thinning up to the critical pinch-off time *t*_*c*_ for all collagen solutions. The breakup of ligaments of the 0.3 and 0.5 mg/mL solutions occurs at the critical pinch-off time *t*_*c*_. The estimated viscous length scale *l*_*sec*_ and viscous time scale *t*_*sec*_ of the secondary microthread for the 0.3 mg/mL solution are 73 nm and 2 ns, respectively. Similarly, the estimated *l*_*sec*_ and *t*_*sec*_ values for the 0.5 mg/mL solution are 120 nm and 5 ns, respectively. Accordingly, a secondary microthread is not observed for the 0.3 and 0.5 mg/mL solutions as the critical pinch-off time *t*_*c*_ approaches. The inertia-controlled ligament pinch-off behavior of the dilute collagen solutions is identical to that of dilute sodium alginate solutions, which was reported by Xu *et al*.^34^ According to those authors, inertial force predominantly resists the capillary thinning of ligaments of dilute sodium alginate solutions (see Herran *et al*.^31^ for overlap concentration of sodium alginate).

**Figure 2.**
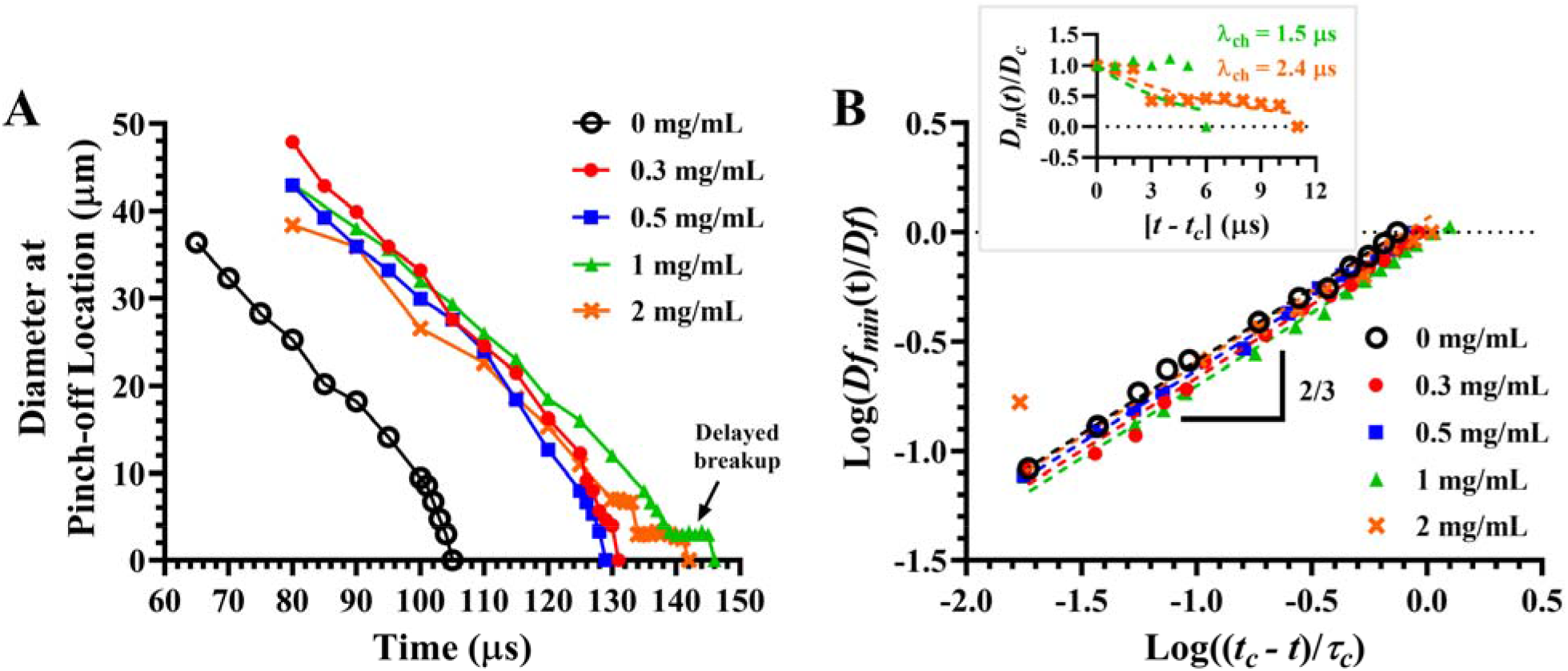
Evolution and breakup of ligaments of collagen solutions with different concentrations, 0 (black open circles), 0.3 (red closed circles), 0.5 (blue squares), 1 (green triangles), and 2 mg/mL (orange crosses). (A) Ligament diameter at the pinch-off location as a function of time. (B) Normalized ligament diameter as a function of normalized shifted time (*t*_*c*_ − *t*). Inset shows the normalized cylindrical microthread diameter as a function of shifted time (*t − t*_*c*_) beyond the critical pinch-off time (*t* > *t*_*c*_) for 1 and 2 mg/mL solutions. Symbols represent experimental values and dashed lines represent model fit values.

The breakup of ligaments of the 1 and 2 mg/mL solutions does not occur at the critical pinch-off time *t*_*c*_. A cylindrical microthread ~ 5 μm in diameter develops at the pinch-off region at the critical pinch-off time *t*_*c*_ for those two solutions (see Figure 1). The microthread for the 1 mg/mL solution lasts for a duration of 5 μs whereas for the 2 mg/mL solution lasts for 10 μs. At the same time, the estimated *l*_*sec*_ and *t*_*sec*_ values for the microthread for the 1 mg/mL solution are 490 nm and 40 ns, respectively. Similarly, the estimated *l*_*sec*_ and *t*_*sec*_ values for the 2 mg/mL solution are 5 μm and 1.4 μs, respectively. The diameter of the microthread decreases exponentially with time for the two solutions as shown in the presented inset of Figure 2. The corresponding characteristic relaxation time *λ*x_*ch*_ for the 1 and 2 mg/mL solution is 1.5 and 2.4 μs, respectively. The estimated viscous length and time scales of the microthread for 1 mg/mL solutions are significantly smaller than the observed length and time scales. Although the estimated viscous length scale of the microthread for 2 mg/mL solution is comparable to the observed length scale, the estimated viscous time scale is much smaller than the observed time scale.

Hence, the elastic force arising because of stretching of the proteins in native conformation and/or denatured proteins in random coil conformation at the solution/air interface of the necked fluid,^20,39,40,48^ predominantly resists the capillary thinning and delays the breakup of the ligaments of 1 and 2 mg/mL solutions beyond the critical pinch-off time *t*_*c*_. This pinch-off behavior is identical to the ligament pinch-off behavior of semidilute unentangled sodium alginate solutions reported by Xu *et al*.^34^ However, the characteristic relaxation time *λ*_*ch*_ of semidilute unentangled sodium alginate solutions is between 36 and 37 μs according to those authors. Thus, in extensional flows, semidilute unentangled collagen solutions show a very weak viscoelastic response when compared to semidilute unentangled sodium alginate solutions. This is because the collagen molecule has a triple helix structure and the three chains which constitute the helix are tightly packed due to the presence of glycine at the center of the helix.^49^ Accordingly, the collagen molecule has a more rigid structure than the sodium alginate molecule (the Mark-Houwink exponent (*a*) value for collagen molecule is 1.8 and for sodium alginate molecule is 0.9).^50,51^

The eventual breakup of ligaments of the 1 and 2 mg/mL solutions is frequently destabilized by aggregates. It is not apparent how aggregates destabilize printing but instabilities generally arise in two ways.^24^ The first is entrapment of air bubbles due to asymmetric meniscus at the orifice, which is caused by presence of aggregates or particles near the orfice. The second is nozzle wetting (which is the accumulation of fluid at the nozzle tip) as shown in Figure 3. The wetting is caused when a microbead forms on the microthread during the elasticity-controlled thinning regime of the ligaments in presence of aggregates. It is not apparent why the satellite bead forms. For example, the microthread breaks up without forming a microbead in some instances (see the image at 155 μs in Figure 3B) whereas it forms a microbead in other instances (see the image at 160 μs in Figure 3B).

**Figure 3.**
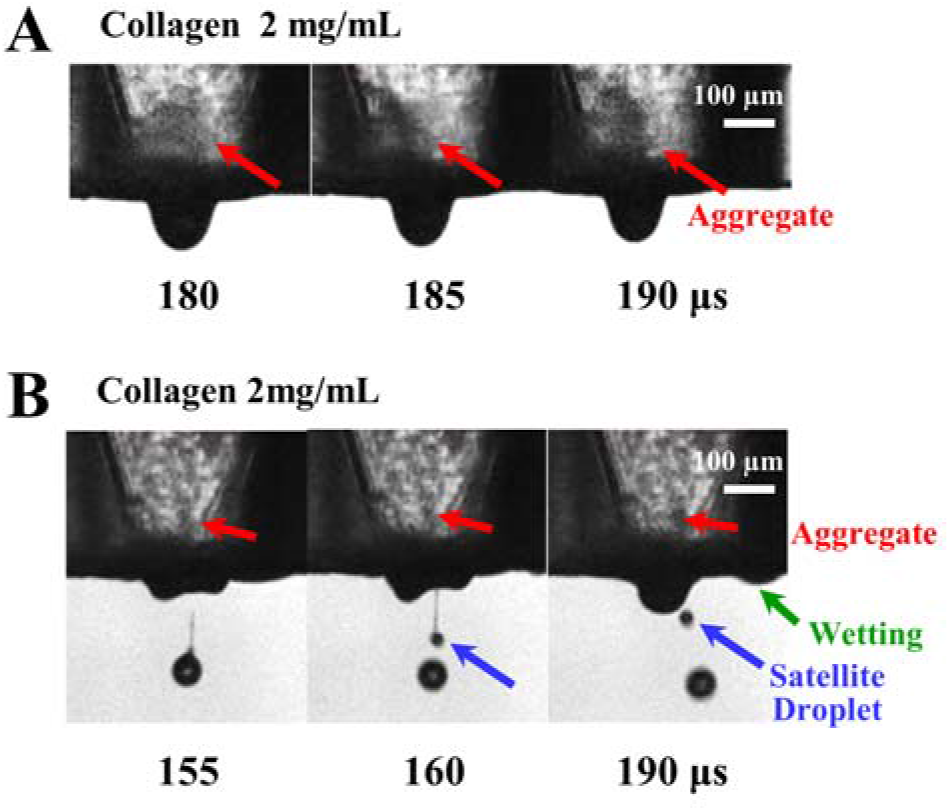
Drop-on-demand inkjet printing of two different samples of collagen solutions with a concentration of 2 mg/mL. (A) The presence of an ~ 150 μm aggregate at the orifice completely stops the formation of droplets. (B) The presence of an ~ 75 μm aggregate at the orifice causes nozzle wetting. The sequential representative images showing single droplet formation, which were assembled by capturing images of different droplets at different time points.

A microbead can form in two ways. The first is when the inertial force dominates viscous, elastic, and capillary forces during the breakup of ligaments.^52^ The second is when the aggregates < 5 μm resist the capillary thinning of the cylindrical microthread and accumulate at one specific region of the microthread.^53^ The microbead eventually breaks up into a satellite droplet having a low kinetic energy, which drifts upwards because of the higher surface energy of the nozzle surface and/or the convection of air towards the nozzle tip due to the downward motion of the primary droplet.^54,55^ The upward drifting satellite droplets, which accumulate at the nozzle orifice, can entrap air bubble during the device actuation. The entrapped air bubbles can dissipate the acoustic pressure waves, transferring insufficient kinetic energy to the fluid near the orifice. Because of the insufficient kinetic energy, formation of droplets can eventually stop. Although aggregates are present in 0.3 and 0.5 mg/mL solutions during printing, they do not destabilize printing.

### Drop-on-demand inkjet printing of surfactant-free fibrinogen solutions

Droplet formation images of dilute fibrinogen solutions with concentrations of 5 mg/mL (*c/c*\* = 0.13), 10 mg/mL (*c/c*\* = 0.26), and 20 mg/mL (*c/c*\* = 0.52) and semidilute unentangled fibrinogen solutions with a concentration of 50 mg/mL (*c/c*\* = 1.30) are presented in Figure 4. Dilute fibrinogen solutions form stable droplets whereas the semidilute unentangled fibrinogen solutions form unstable droplets. The dwell time for the solutions is 20 μs, which is identical to dwell time for Milli-Q water. According to eq 1, the dwell time for the high energy droplet formation with the maximum possible ejection velocity for Milli-Q water is 21 μs. The ligaments of the solutions emerge out of the orifice at 60 μs, which is similar to Milli-Q water. Droplets of dilute fibrinogen solutions pinch off at 110 μs, which is comparable to Milli-Q water. However, droplets of 50 mg/mL solutions pinch-off from the ligaments at 160 us, which is 50 μs longer when compared to Milli-Q water. This longer pinch-off time is because of viscoelasticity and a lower back pressure of −10 mmHg.

**Figure 4.**
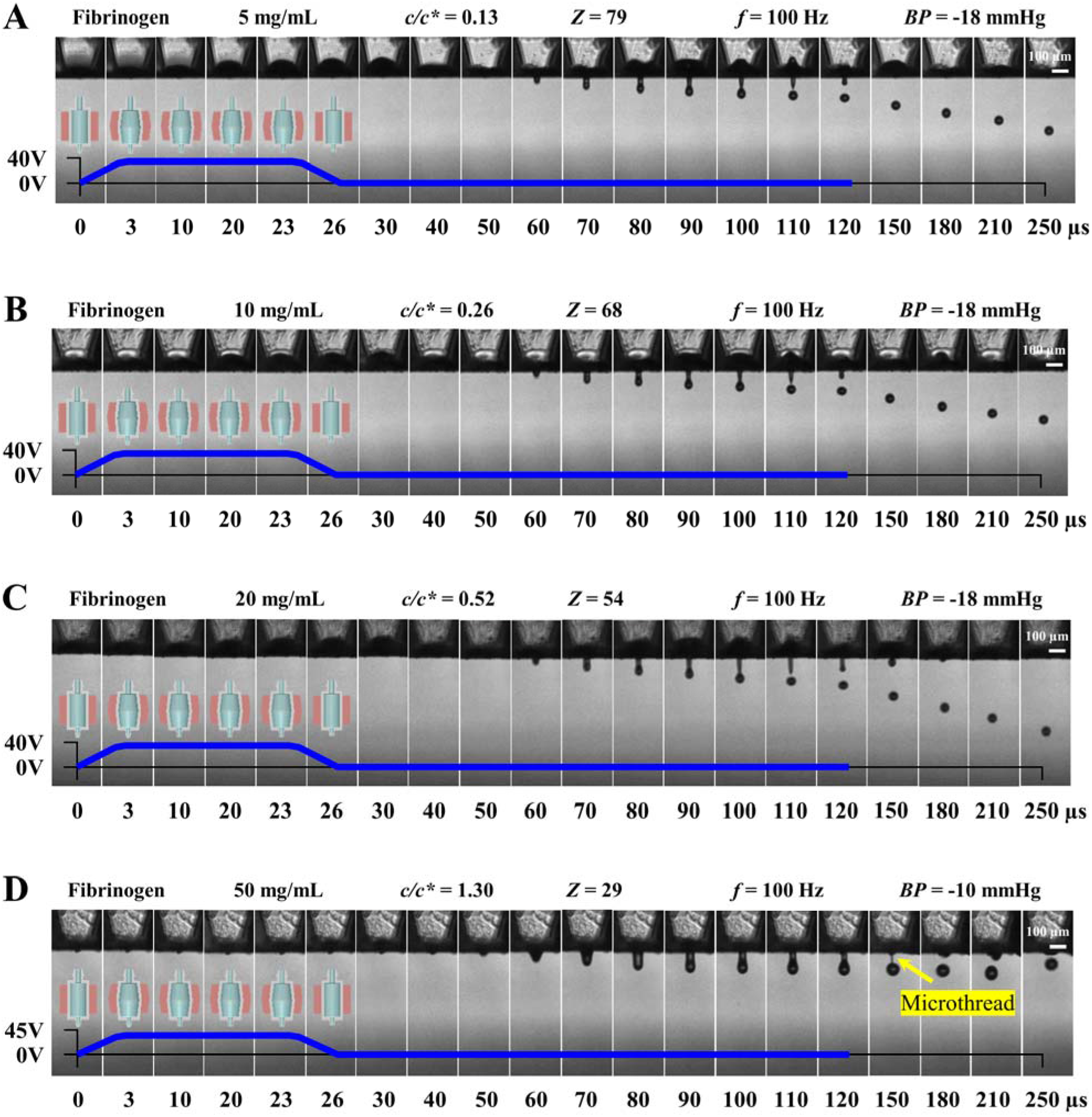
Drop-on-demand inkjet printing of dilute and semidilute unentangled fibrinogen solutions. (A) 5 mg/mL, (B) 10 mg/mL, (C) 20 mg/mL, and (D) 50 mg/mL. The *Z* values are based on the nozzle orifice diameter. The sequential representative images showing single droplet formation, which were assembled by capturing images of different droplets at different time points.

The diameter of ligaments at the pinch-off region as function of time is presented in Figure 5. The minimum ligament diameter *D_f_min__*(*t*) scales as (*t*_*c*_ – *t*)/τ_*c*_ to the power of 2/3 up to the critical pinch-off time *t*_*c*_ for dilute and semidilute unentangled fibrinogen solutions. Accordingly, inertial force predominantly resists the capillary thinning up to the critical pinch-off time *t*_*c*_ for all those solutions. The breakup of ligaments of the 5, 10, and 20 mg/mL solutions occurs at the critical pinch-off time *t*_*c*_. The estimated values of the viscous length scale *l*_*sec*_ for those solutions are 19, 26, and 41 nm, respectively. Similarly, the estimated values of the viscous time scale *t*_*sec*_ for those solutions are 0.4, 0.6, and 1.2 ns, respectively. Accordingly, a secondary microthread is not observed for those solutions as the critical pinch-off time *t*_*c*_ approaches. This behavior is identical to the ligament pinch-off behavior of dilute collagen solutions.

**Figure 5.**
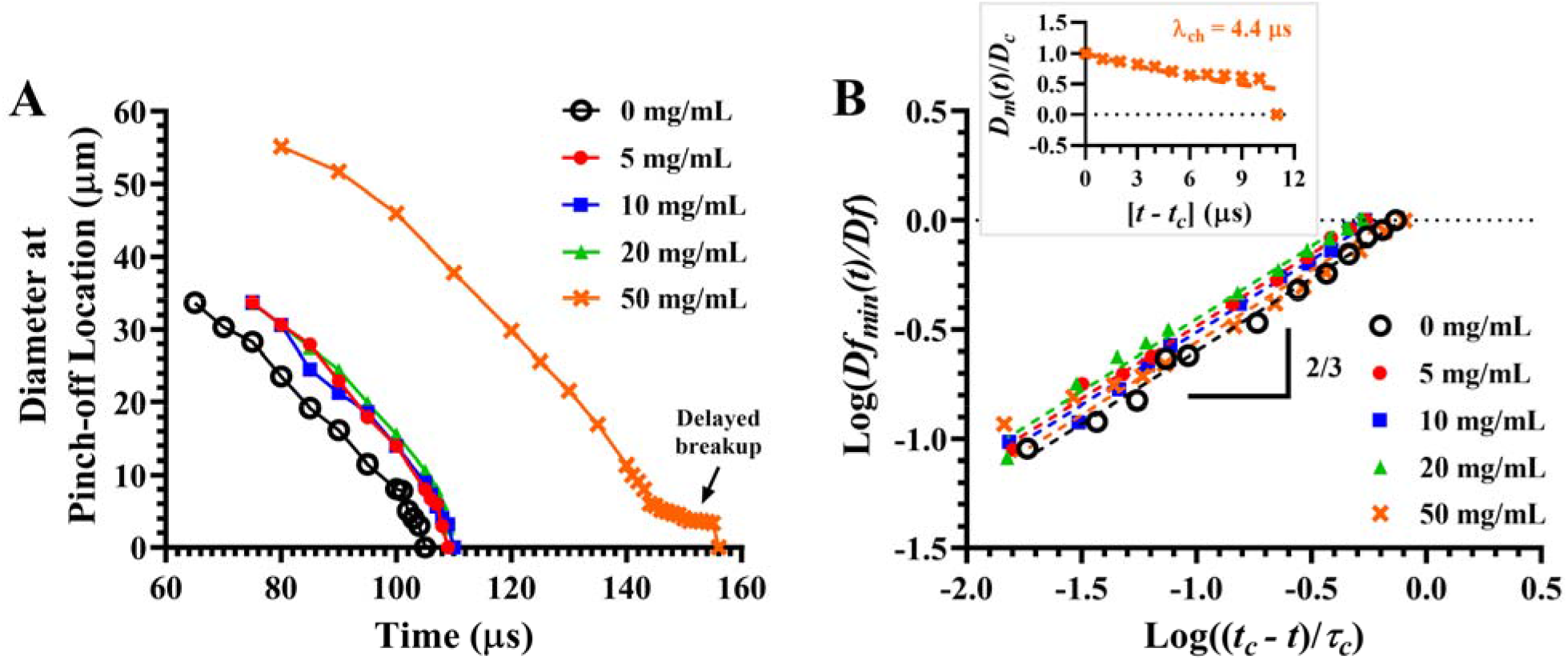
Evolution and breakup of ligaments of fibrinogen solutions with different concentrations, 0 (black open circles), 5 (red closed circles), 10 (blue squares), 20 (green triangles), and 50 mg/mL (orange crosses). (A) Ligament diameter at the pinch-off location as a function of time. (B) Normalized ligament diameter as a function of normalized shifted time (*t*_*c*_ − *t*). Inset shows the normalized cylindrical microthread diameter as a function of shifted time (*t - t*_*c*_) beyond the critical pinch-off time (*t > t*_*c*_) for 50 mg/mL solution. Symbols represent experimental values and dashed lines represent model fit values.

The breakup of ligaments of the 50 mg/mL solutions does not occur at the critical pinch-off time *t*_*c*_ and a cylindrical microthread of ~ 5 μm in diameter develops at the pinch-off region at the critical pinch-off time *t*_*c*_ (see Figure 4). The microthread lasts for a duration of 11 μs, which is interestingly comparable to that of much viscous 2 mg/mL collagen solutions. The estimated viscous length scale *l*_*sec*_ and the viscous time scale *t*_*sec*_ of the microthread are 140 nm and 8 ns, respectively. At the same time, the diameter of the microthread decreases exponentially with time and the corresponding characteristic relaxation time *λ*_*ch*_ is 4.4 μs as shown in Figure 5. This value is longer than the characteristic relaxation time of the semidilute unentangled collagen solutions because of the flexible structure of fibrinogen molecules.^27^ Because the viscous length and time scales of the microthread are significantly smaller than the observed length and time scales, the elastic force predominantly resists the capillary thinning and delays the breakup of the ligaments for 50 mg/mL solutions beyond the critical pinch-off time *t*_*c*_, similar to the semidilute collagen solutions. The eventual breakup of the ligaments of those semidilute unentangled fibrinogen solutions is frequently destabilized by aggregates, similar to semidilute untangled collagen solutions. Although aggregates are observed in dilute fibrinogen solutions during printing (see Figure S8 in the Supporting Information), they do not destabilize the formation of droplets. Aggregates as big as 100 μm are frequently observed during the printing of 50 mg/mL solutions, which destabilize the formation of droplets. Those aggregates, when present near the orifice, disrupt the printing in a way similar to semidilute unentangled collagen solutions (see Figure 6, i.e., nozzle wetting and/or asymmetric meniscus, which can entrap air).

**Figure 6.**
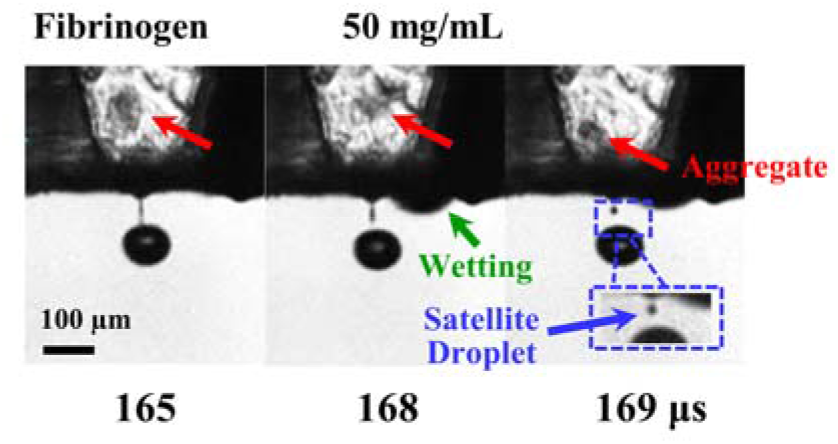
Drop-on-demand inkjet printing of fibrinogen solution with a concentration of 50 mg/mL. The presence of an ~ 100 μm aggregate at the orifice causes nozzle wetting. The sequential representative images showing single droplet formation, which were assembled by capturing images of different droplets at different time points.

### Drop-on-demand inkjet printing of surfactant-free thrombin solutions

Droplet formation images of 20 U/mL thrombin solutions and the corresponding actuation voltage pulse properties are presented in Figure 7 (see Figure S9 in the Supporting Information for droplet formation images of 5, 10, and 15 U/mL thrombin solutions). The solutions form stable droplets. In general, thrombin solutions with a concentration *c* < 1000 U/mL are very dilute (*c/c*\* ≪ 1) and their behavior is analogous to Milli-Q water during inkjet printing. Accordingly, the actuation voltage pulse properties for the solutions are identical to those of Milli-Q water. Further, the ligaments of the solutions emerge out of the orifice at 60 μs and droplets pinch off from them at 105 – 110 μs, which is similar to Milli-Q water.

**Figure 7.**
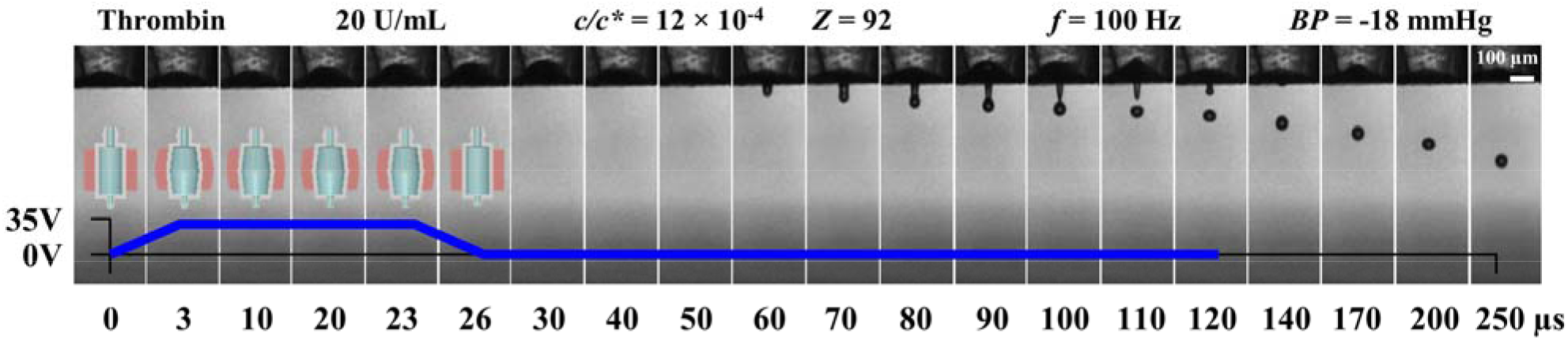
Drop-on-demand inkjet printing of thrombin solutions with a concentration of 20 U/mL. The *Z* value are based on the nozzle orifice diameter. The sequential representative images showing single droplet formation, which were assembled by capturing images of different droplets at different time points.

The diameter of the ligaments at the pinch-off region as function of time is presented in Figure 8. The minimum ligament diameter *D_f_min__*(*t*) scales as (*t*_*c*_ – *t*)/τ_*c*_ to the power of 2/3 up to the critical pinch-off time *t*_*c*_. Hence, inertial force predominantly resists the capillary thinning up to the critical pinch-off time *t*_*c*_. The breakup of the ligaments occurs at the critical pinch-off time *t*_*c*_. The estimated viscous length scale *l*_*sec*_ of the solutions is 11 – 14 ns whereas the estimated viscous time scale *t*_*sec*_ is 0.1 – 0.3 ns. Hence, a secondary microthread is not observable for the solutions (see Figure 7) as the critical pinch-off time *t*_*c*_ approaches. Although aggregates are present in the solutions, they do not disrupt printing.

**Figure 8.**
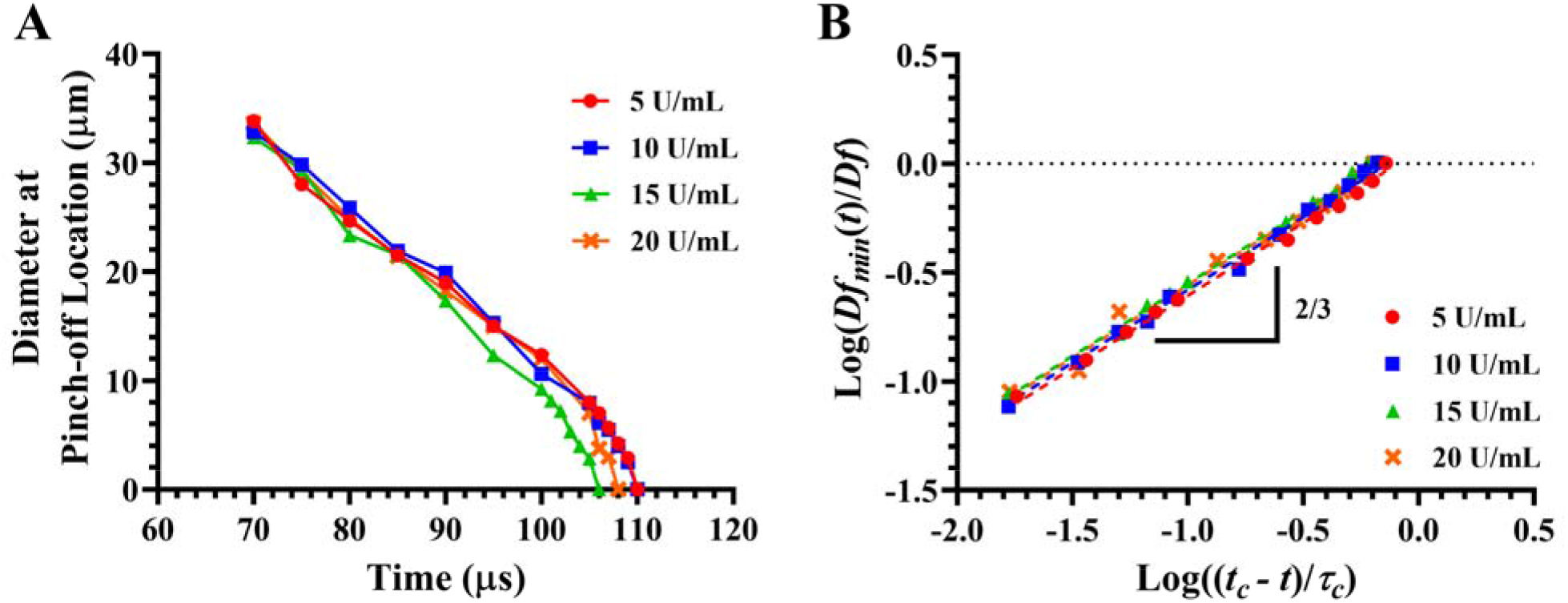
Evolution and breakup of ligaments of thrombin solutions with different concentrations, 5 (red circles), 10 (blue squares), 15 (green triangles), and 20 U/mL (orange crosses). (A) Ligament diameter at the pinch-off location as a function of time. (B) Normalized ligament diameter as a function of normalized shifted time (*t_c_ - t*).

### Drop-on-demand inkjet printing of filtered-surfactant-free protein solutions

Collagen solutions with concentrations of 1 and 2 mg/mL, and fibrinogen solutions with concentrations of 10 and 50 mg/mL were filtered just before printing to remove aggregates, which might have formed during storage or during sample preparation because of the bulk aggregation of proteins. The length of collagen monomers is 300 nm and filtering of collagen solutions with a 450 nm (0.45 μm) pore size syringe filter causes considerable protein loss.^56^ In this work, the loss was ~ 20 % of the solution volume for the 1 mg/mL solutions and ~ 40 % of the solution volume for the 2 mg/mL solutions because of filter clogging. It is also known that, the length of fibrinogen monomers is 46 nm and filtering of fibrinogen solutions with a 220 nm (0.22 μm) pore size syringe filter does not cause significant loss of the protein.^57^

During printing of unfiltered 10 mg/mL fibrinogen solutions, aggregates were observed near the orifice, which were momentarily decreasing the velocity of ejected droplets as shown in Figure 9A. This is reflected by a standard deviation of 0.032 m/s in the presented velocity plot (calculated based on the relative position of droplets with respect to the nozzle orifice at 500 μs after actuation) of Figure 9B. Filtering of those solutions improved droplet formation, decreasing the standard deviation to 0.017 m/s, which is perhaps because of the absence of aggregates. However, the standard deviation for Milli-Q water droplets is 0.010 m/s, which signifies the presence of aggregates despite the filtering of 10 mg/mL fibrinogen solutions. Similarly, filtering of 1 mg/mL collagen solutions and 50 mg/mL fibrinogen solutions improved droplet formation (see Figure S10 in the Supporting Information). However, filtering of 2 mg/mL collagen solutions did not improve droplet formation. Aggregates were present during printing despite the filtering of 2 mg/mL collagen and 50 mg/mL fibrinogen solutions as shown in Figs. 9 C-D. Those aggregates prevent the identification of optimal dwell time for 2 mg/mL collagen solutions because of inconsistent droplet formation (see Figure S10 in the Supporting Information). Hence, aggregates are not only forming during storage and preparation of samples but also during inkjet printing. Note that aggregates were observed during the printing of freshly prepared (1 – 2 days old) unfiltered fibrinogen solutions.

**Figure 9.**
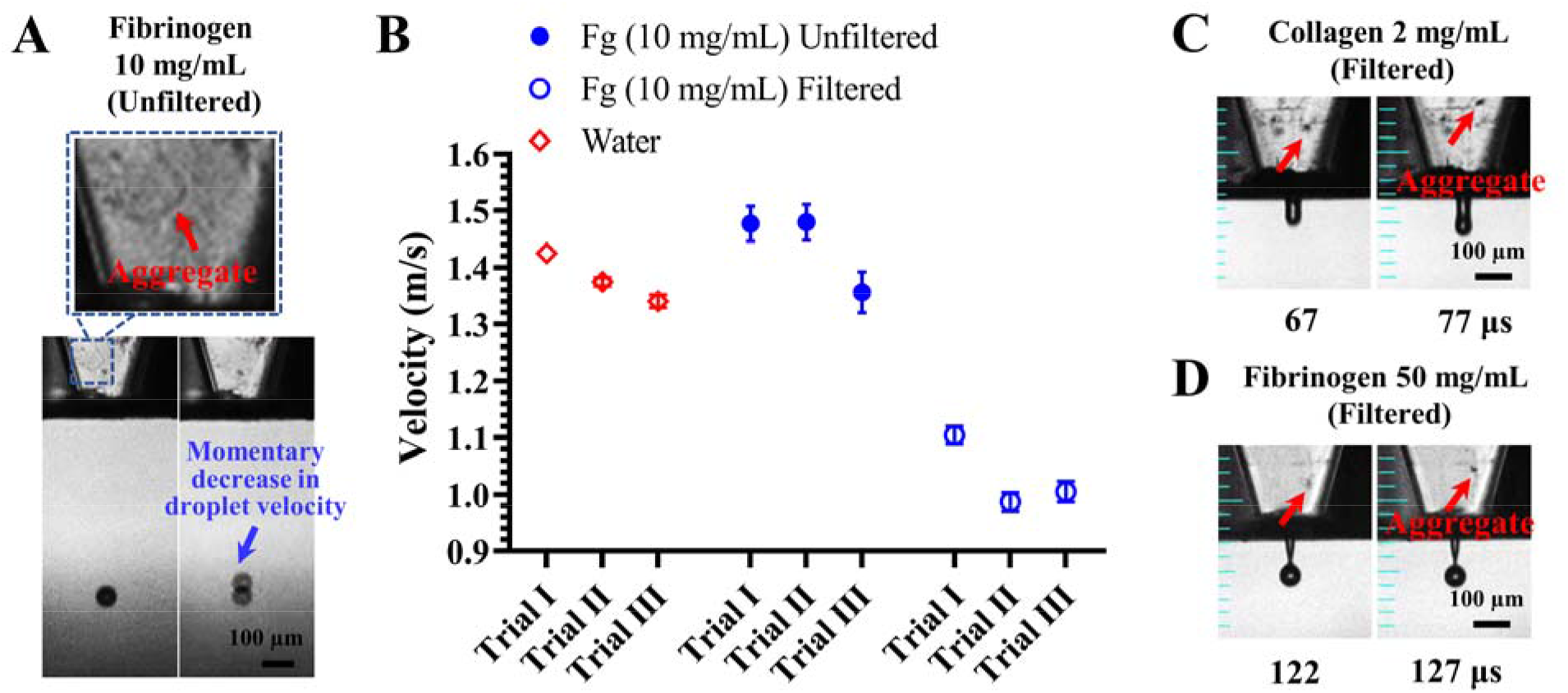
Formation of aggregates in unfiltered and filtered protein solutions during inkjet printing. (A) Unfiltered fibrinogen solution with a concentration of 10 mg/mL and an aggregate near the orifice momentarily decreases droplet velocity. (B) Droplet velocity of unfiltered (blue closed circles) and filtered (blue open circles) fibrinogen solutions with a concentration of 10 mg/mL and Milli-Q water (red open diamonds) from different trials. Mean data and standard deviation are plotted (*n* = 30 replicates). (C) Filtered collagen solution with a concentration of 2 mg/mL. (D) Filtered fibrinogen solution with a concentration of 50 mg/mL.

## Conclusions

Protein concentration controls droplet formation and breakup of surfactant-free collagen, fibrinogen, and thrombin solutions during inkjet printing. The capillary thinning of ligaments of dilute collagen, fibrinogen, and thrombin solutions solutions is predominantly resisted by inertial force up to the break up point. The capillary breakup of ligaments of semidilute unentangled collagen and fibrinogen solutions is predominantly resisted by inertial force on the initial onset of necking. However, the elastic force delays the breakup of those solutions. The resistance of viscous force to the capillary thinning of the dilute and semidilute unentangled protein solutions is negligible. The dilute protein solutions form stable droplets, employing a simple unipolar trapezoidal voltage pulse. However, the formation of stable droplets of semidilute unentangled collagen and fibrinogen solutions is constantly disrupted by the presence of aggregates or subvisible particles. Although aggregates are present in dilute protein solutions, they do not disrupt bioprinting.

In view of these results, long term storage of protein solutions is not recommended for the formation of stable droplets during inkjet bioprinting of the surfactant-free collagen, fibrinogen, and thrombin solutions. In addition, filtering of those protein solutions is recommended when it does not cause significant loss of proteins. Further, successful delivery of small volumes of semidilute unentangled collagen and fibrinogen solutions requires novel materials and novel inkjet nozzle or printer design strategies. Such materials minimize the protein adsorption at the solution/solid interface and such strategies minimize the solution/air and solution/solid interfaces of protein solutions during inkjet bioprinting.

## Associated Content

### Supporting Information

Drop-on-demand inkjet printing of water and surfactant-free collagen, fibrinogen, and thrombin solutions. Droplet/substrate interactions of impinging droplets on a smooth solid surface. Dimensionless parameters and time scales associated with breakup of droplets. Dimensionless parameters associated with impinging droplets.

## Author Information

### Notes

The authors declare no competing financial interest.

## Acknowledgements

The authors are grateful for the valuable insights provided by Dr. Ralph H. Colby from the Department of Materials Science and Engineering at Penn State on the behavior of protein molecules in an extensional flow and at the liquid/air and liquid/solid interfaces. The authors are grateful for the guidance of Dr. Bruce Gluckman from the Engineering Science and Mechanics Department at Penn State on the planning of experiments. This research has been partially supported by the Osteology Foundation Award #15-042 and the Hartz Career Development Professorship (I.T.O.).

---

For Table of Contents Only

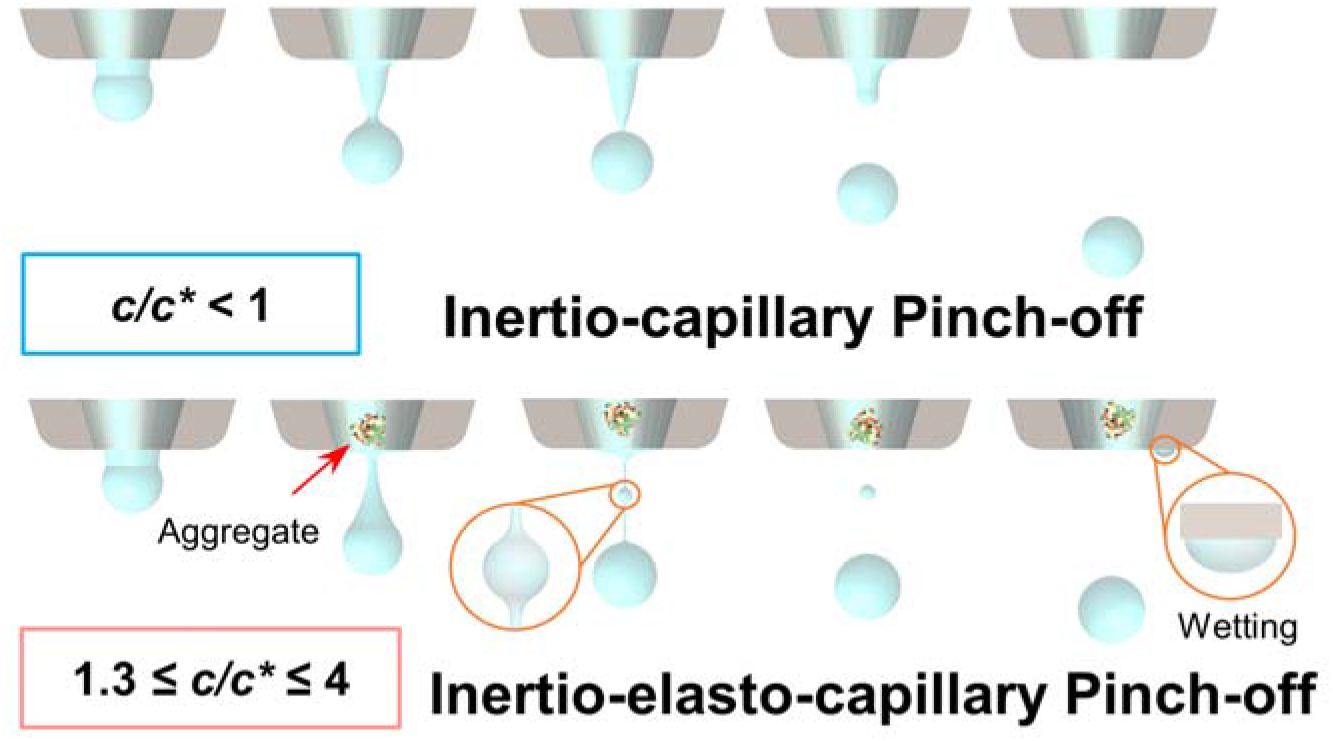

