## Supplementary Information for "The role of concentration on drop formation and breakup of collagen, fibrinogen, and thrombin solutions during inkjet bioprinting"

### Materials and methods

#### Statistical analysis

In order to compare the diameter and velocity of water droplets formed with different back pressure, dwell time, rise/fall times, and frequency, two-sided one-way ANOVA with posthoc Fisher's individual tests (assuming equal variances) was used at a 95% confidence level in order to determine the significance levels. Differences were considered significant at $p^{*} < 0.05$, $p^{**} < 0.01$, and $p^{***} < 0.001$. All statistical analysis was performed using Minitab 17.3 (Minitab Inc.).

#### Analysis of droplet/substrate interactions

Droplets of protein solutions were deposited on 18 × 18 mm micro cover glass slides (cat. no. 4833045, VWR International, West Chester, PA) to check splashing. The droplets were deposited on the glass slides placed at distance of 1.2 – 1.5 mm below the nozzle tip while moving the nozzle with a horizontal velocity of 1 mm/s.

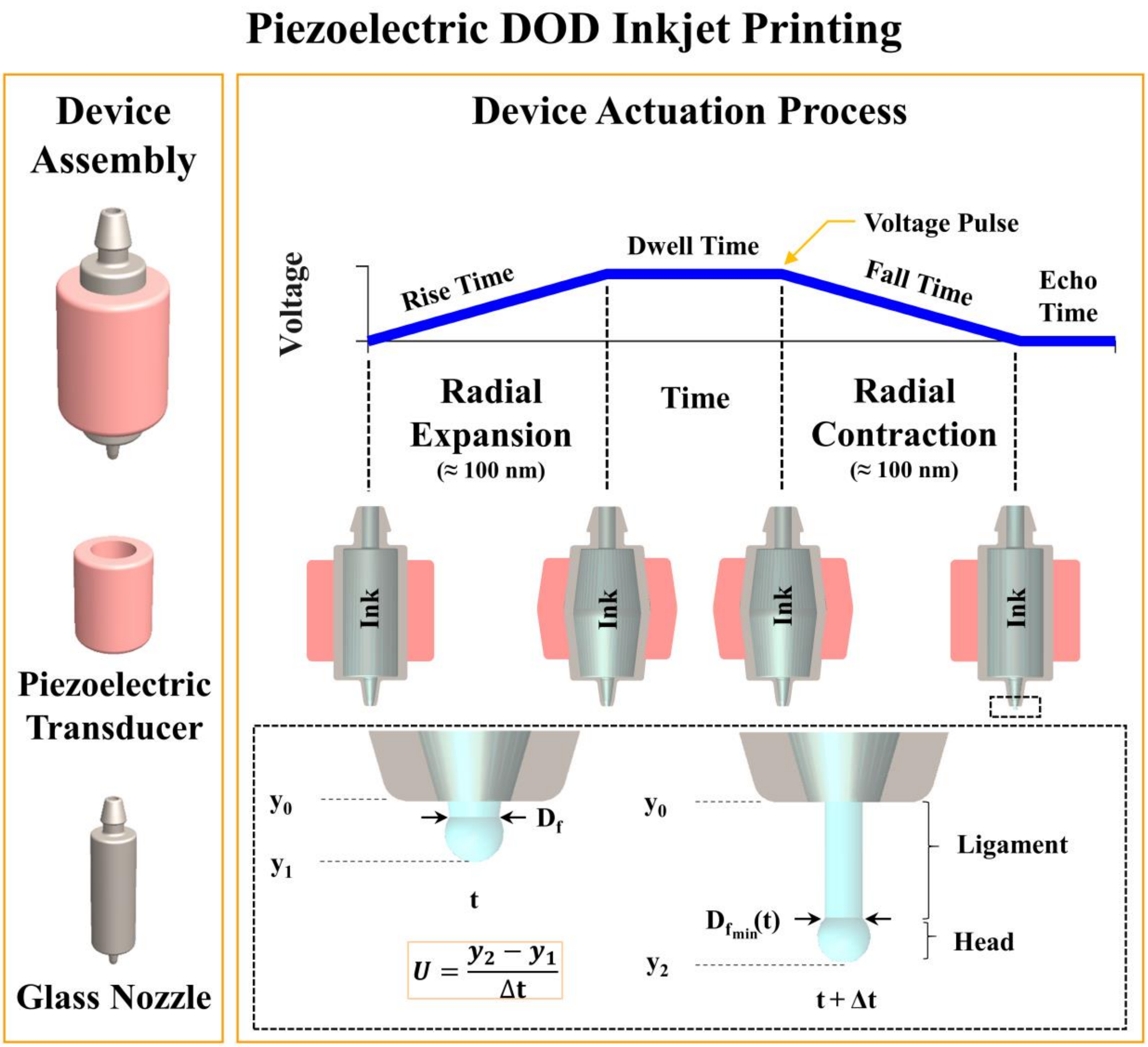

**Figure S1.** Piezoelectric DOD inkjet dispensing device and its actuation with a trapezoidal voltage pulse.

### Results and discussion

#### Drop-on-demand inkjet printing of Milli-Q water

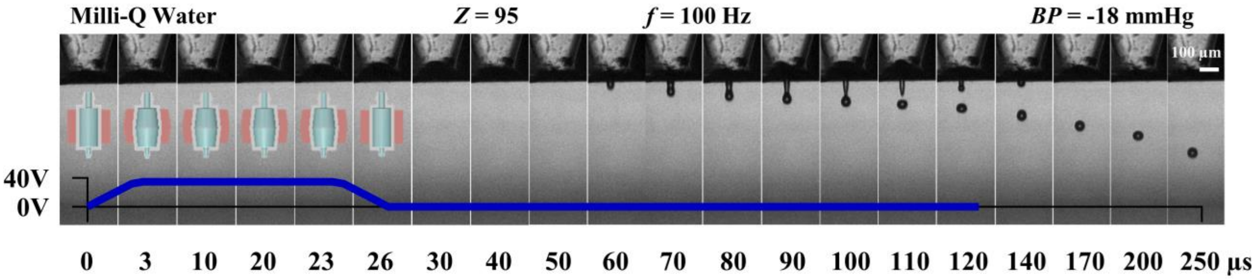

**Figure S2.** Drop-on-demand inkjet printing of Milli-Q water. The sequential representative images showing single droplet formation, which were assembled by capturing images of different droplets at different time points.

Water forms stable droplets with a back pressure of -18 mmHg and a unipolar trapezoidal voltage pulse having a rise time ($t_{rise}$) of 3 µs, dwell time ($t_{dwell}$) of 20 µs, fall time ($t_{fall}$) of 3 µs, echo time ($t_{echo}$) of 100 µs, dwell voltage ($V_{dwell}$) of 40 V, and frequency of 100 Hz. This experimental dwell time is comparable to the theoretical optimal dwell time of 21 µs for the high energy droplet formation with the maximum possible ejection velocity, which is determined with the relationship, $t_{dwell} = l/C$. The capillary length $l$ = 30 mm and $C$ =1450 m/s in water.[1]

A ligament emerges out of the orifice at 60 µs and a droplet pinches off from it at 110 µs (i.e., critical pinch-off time $t_c$). The minimum ligament diameter $D_{f_{min}}(t)$ scales as $(t_c - t)/\tau_c$ to the power of 2/3 up to the critical pinch-off time $t_c$, regardless of the operating conditions (see Figure S3). Hence, the inertial force predominantly resists capillary thinning of the ligament up to critical pinch-off time $t_c$. At $t_c$, the estimated viscous length scale $l_{sec}$ and viscous time scale $t_{sec}$ of the secondary microthread are 10 nm and 0.2 ns, respectively. Accordingly, a secondary microthread is not observed and the breakup of the ligament occurs at the critical pinch-off time $t_c$.

The diameter and velocity of the ligament are controlled by back pressure, rise time, dwell time, and fall time (see Figures S4-7). In general, back-pressure controls the position and shape of the meniscus, controlling the kinetic energy distributed to the water near the nozzle orifice.[2,3] Whereas, the dwell time controls the time at which the piezoelectric transducer radially deforms to dampen the dilation pressure wave. [4,5] Rise and fall times control the amplitude of the acoustic pressure wave, which control the diameter and velocity of water droplets.[5] A frequency above 200 Hz was destabilizing the formation of water droplets perhaps because of undissipated residual acoustic pressure waves, which requires further investigation.

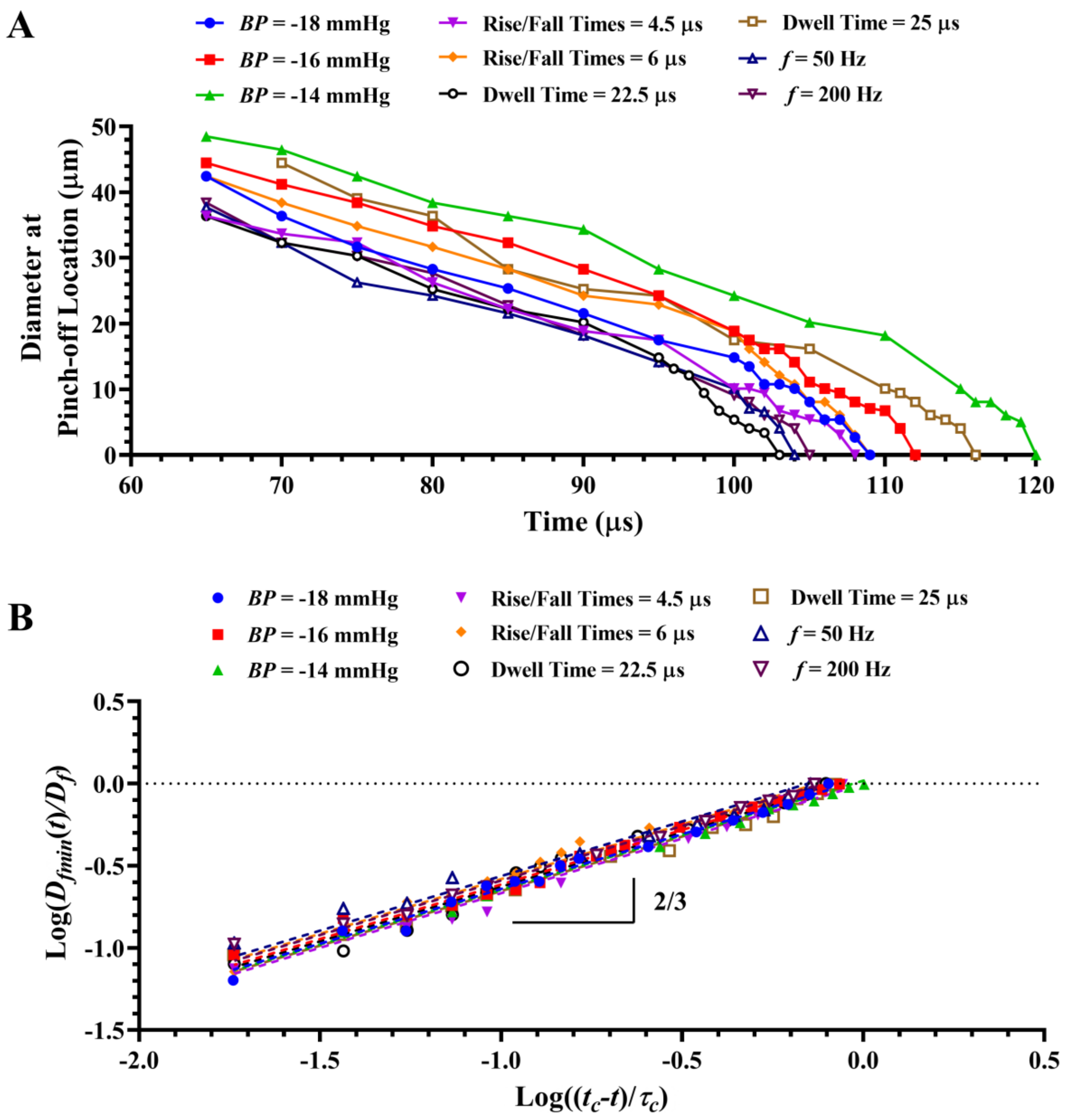

**Figure S3.** Evolution and breakup of ligaments of Milli-Q water under different conditions. The blue color solid circles represent a back pressure (BP) of -18 mmHg, rise/fall times of 3 μs, dwell time of 20 μs, echo time of 100 μs, dwell voltage of 40 V, and a frequency of 100 Hz. Other symbols represent a particular condition, such as back pressure, rise/fall times, dwell time, or frequency, which was changed. Ligament diameter variation at front-pinching location as a function of time under different conditions. (C) Normalized ligament diameter as a function of normalized shifted time ($t_c$-$t$) under different conditions.

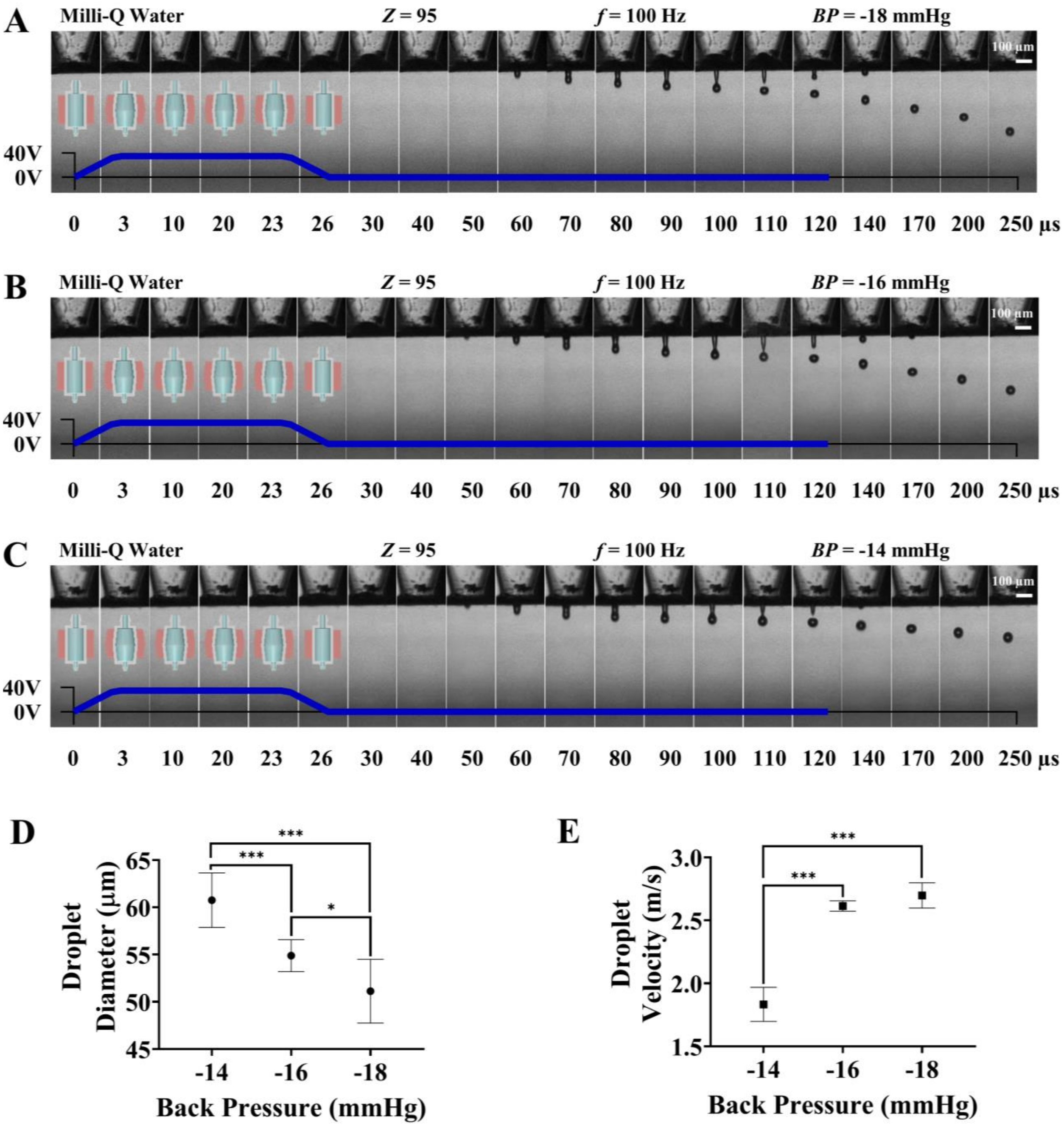

**Figure S4.** Drop-on-demand inkjet printing of Milli-Q water with different back pressures. (A) -18 mmHg, (B) -16 mmHg, and (C) -14 mmHg. The sequential representative images showing single droplet formation, which were assembled by capturing images of different droplets at different time points. (D) Droplet diameter as a function of back pressure (*n* = 9 replicates). (E) Droplet velocity as a function of back pressure (*n* = 3 replicates).

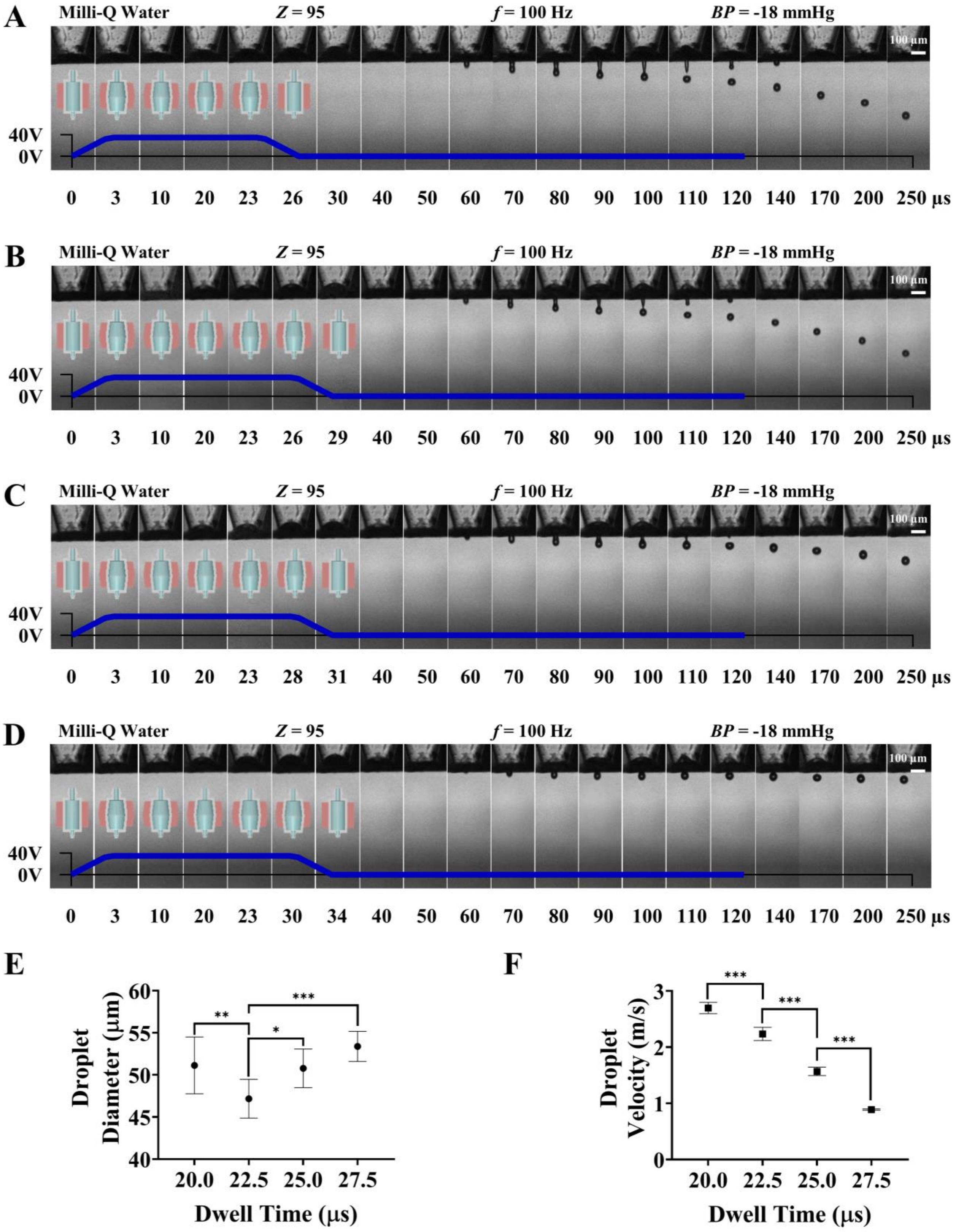

A
Milli-Q Water
Z = 95
f = 100 Hz
BP = -18 mmHg
100 µm
40V
0V
0 3 10 20 23 26 30 40 50 60 70 80 90 100 110 120 140 170 200 250 µs
B
Milli-Q Water
Z = 95
f = 100 Hz
BP = -18 mmHg
100 µm
40V
0V
0 3 10 20 23 26 29 40 50 60 70 80 90 100 110 120 140 170 200 250 µs
C
Milli-Q Water
Z = 95
f = 100 Hz
BP = -18 mmHg
100 µm
40V
0V
0 3 10 20 23 28 31 40 50 60 70 80 90 100 110 120 140 170 200 250 µs
D
Milli-Q Water
Z = 95
f = 100 Hz
BP = -18 mmHg
100 µm
40V
0V
0 3 10 20 23 30 34 40 50 60 70 80 90 100 110 120 140 170 200 250 µs
E
Droplet Diameter (µm)
40 45 50 55 60
**
*
***
20.0 22.5 25.0 27.5
Dwell Time (µs)
F
Droplet Velocity (m/s)
0 1 2 3
***
***
***
20.0 22.5 25.0 27.5
Dwell Time (µs)

**Figure S5.** Drop-on-demand inkjet printing of Milli-Q water with different dwell times. (A) 20 μs, (B) 22.5 μs, (C) 25 μs, and (D) 27.5 μs. The sequential representative images showing single droplet formation, which were assembled by capturing images of different droplets at different time points. (E) Droplet diameter as a function of dwell time (*n* = 9 replicates). (F) Droplet velocity as a function of dwell time (*n* = 3 replicates).

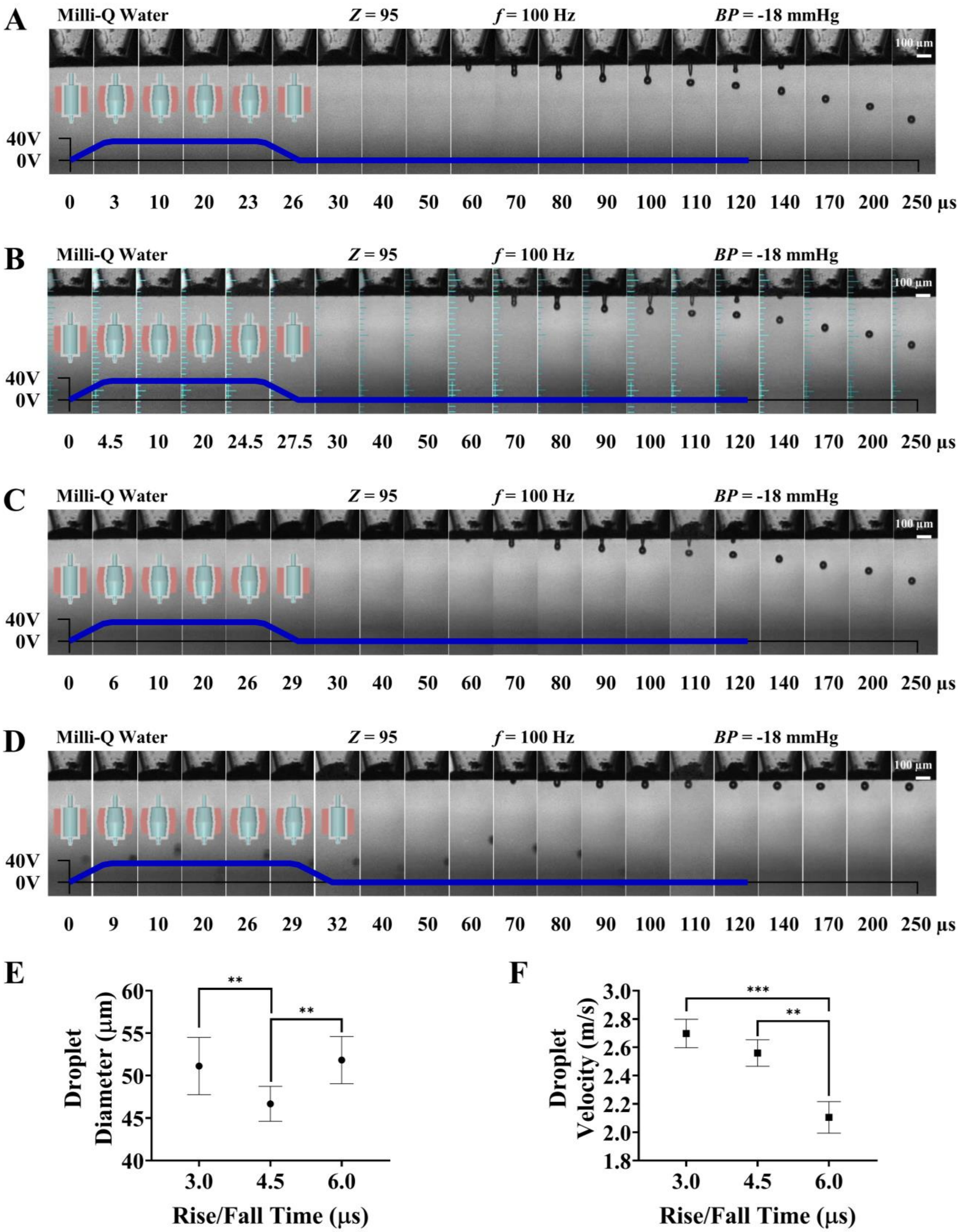
A
Milli-Q Water
Z = 95
f = 100 Hz
BP = -18 mmHg
100 µm
40V
0V
0 3 10 20 23 26 30 40 50 60 70 80 90 100 110 120 140 170 200 250 µs
B
Milli-Q Water
Z = 95
f = 100 Hz
BP = -18 mmHg
100 µm
40V
0V
0 4.5 10 20 24.5 27.5 30 40 50 60 70 80 90 100 110 120 140 170 200 250 µs
C
Milli-Q Water
Z = 95
f = 100 Hz
BP = -18 mmHg
100 µm
40V
0V
0 6 10 20 26 29 30 40 50 60 70 80 90 100 110 120 140 170 200 250 µs
D
Milli-Q Water
Z = 95
f = 100 Hz
BP = -18 mmHg
100 µm
40V
0V
0 9 10 20 26 29 32 40 50 60 70 80 90 100 110 120 140 170 200 250 µs
E
Droplet Diameter (µm)
60
55
50
45
40
**
**
3.0 4.5 6.0
Rise/Fall Time (µs)
F
Droplet Velocity (m/s)
3.0
2.8
2.6
2.4
2.2
2.0
1.8
***
**
3.0 4.5 6.0
Rise/Fall Time (µs)

**Figure S6.** Drop-on-demand inkjet printing of Milli-Q water with different rise/fall times. (A) 3 μs, (B) 4.5 μs, (C) 6 μs, and (D) 9 μs. The sequential representative images showing single droplet formation, which were assembled by capturing images of different droplets at different time points. (E) Droplet diameter as a function of rise/fall times (*n* = 9 replicates). (F) Droplet velocity as a function of rise/fall times (*n* = 3 replicates).

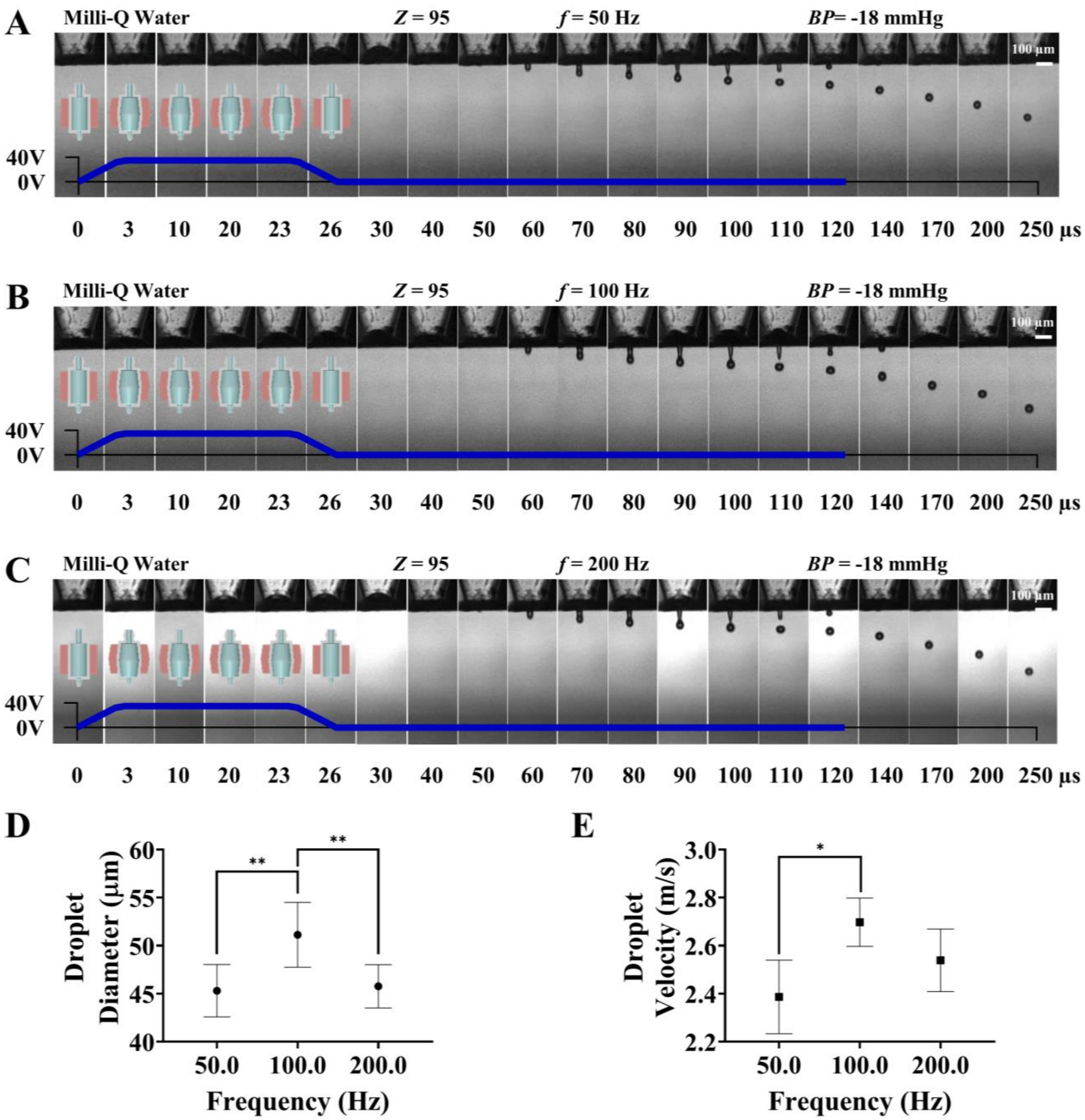

**Figure S7.** Drop-on-demand inkjet printing of Milli-Q water with different frequencies. (A) 50 Hz, (B) 100 Hz, and (C) 200 Hz. The sequential representative images showing single droplet formation, which were assembled by capturing images of different droplets at different time points. (D) Droplet diameter as a function of frequency (*n* = 9 replicates). (E) Droplet velocity as a function of frequency (*n* = 3 replicates).

#### Drop-on-demand inkjet printing of surfactant-free fibrinogen solutions

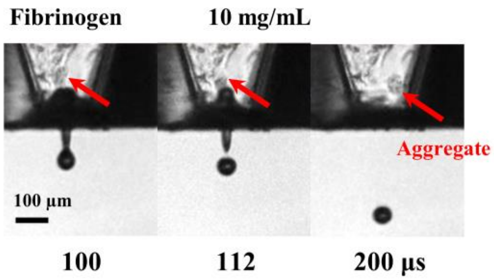

**Figure S8.** Drop-on-demand inkjet printing of fibrinogen solution with a concentration of 10 mg/mL. The presence of an ~ 40 µm aggregate at the orifice does not disrupt droplet formation. The sequential representative images showing single droplet formation, which were assembled by capturing images of different droplets at different time points.

**Drop-on-demand inkjet printing of surfactant-free thrombin solutions**

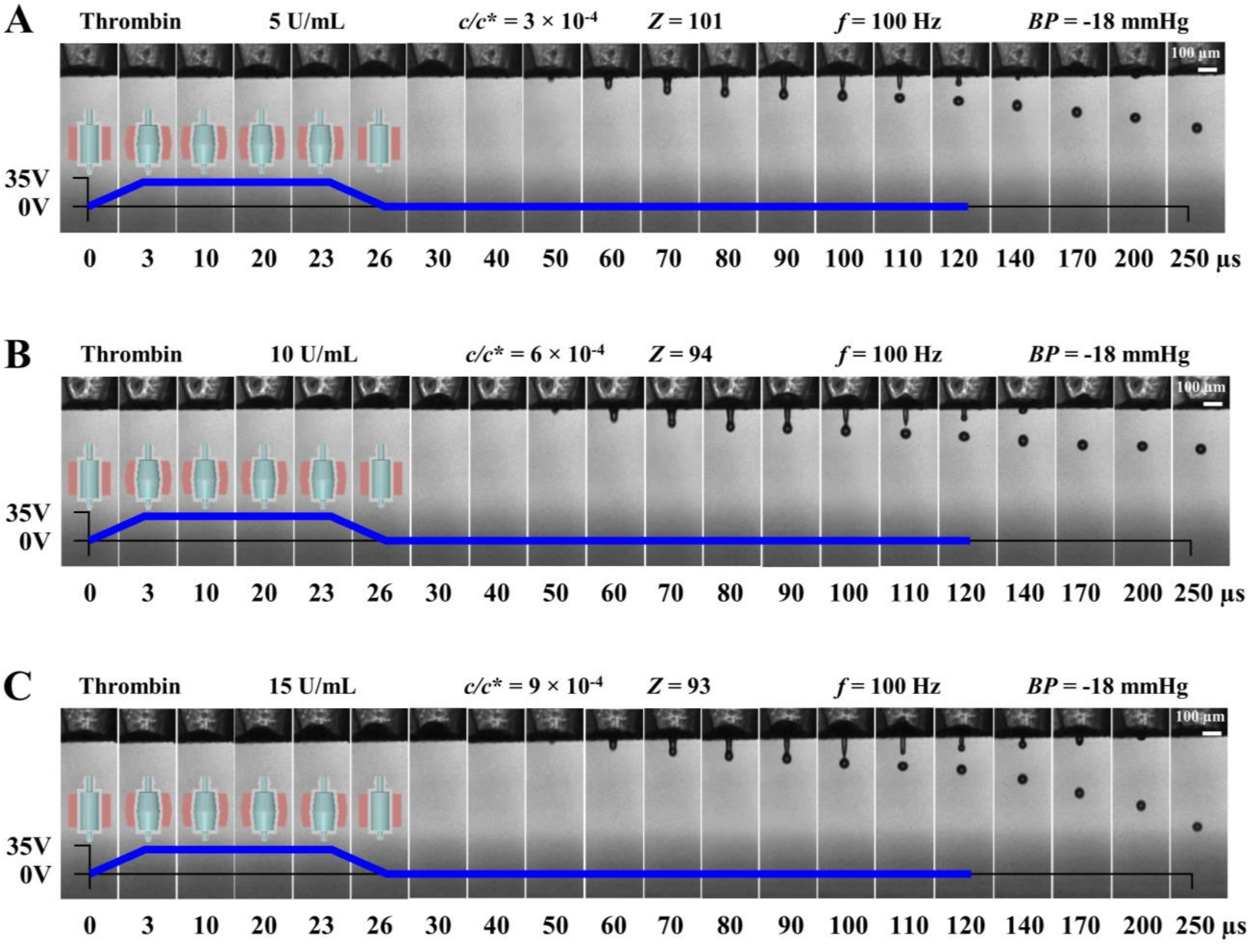

**Figure S9.** Drop-on-demand inkjet printing of thrombin solutions with different concentrations. (A) 5 U/mL, (B) 10 U/mL, and (C) 15 U/mL. The *Z* values are based on the nozzle orifice diameter. The sequential representative images showing single droplet formation, which were assembled by capturing images of different droplets at different time points.

**Drop-on-demand inkjet printing of filtered collagen and fibrinogen solutions**

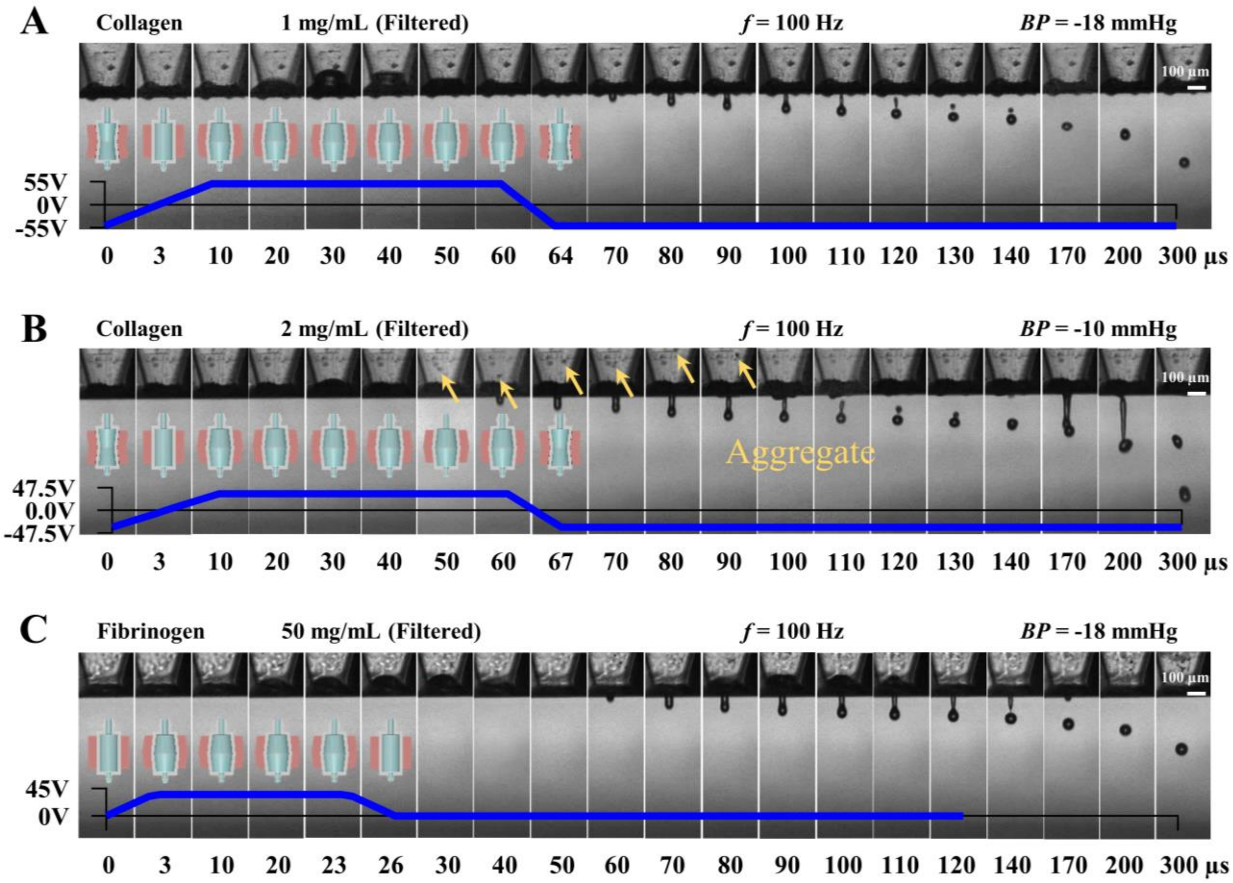

**Figure S10.** Drop-on-demand inkjet printing of semidilute untangled protein solutions which were filtered prior to printing. (A) 1 mg/mL collagen solution filtered with a 0.45 µm pore size filter. (B) 2 mg/mL collagen solution filtered with a 0.45 µm pore size filter. (C) 50 mg/mL fibrinogen solution filtered with a 0.22 µm pore size filter. The sequential representative images showing single droplet formation, which were assembled by capturing images of different droplets at different time points.

#### Droplet/substrate interactions

Polymerization (or crosslinking) of collagen and fibrinogen solutions occurs post deposition on a substrate. Collagen solutions at pH 7 polymerize within 5 – 15 min at room temperature. Similarly, fibrinogen solutions polymerize into fibrin through the enzymatic action of thrombin within 5 – 15 min at room temperature. Schiaffino *et al*.[6] studied the spreading behavior of molten droplets with Weber (*We*) number and the initial stages during the spreading of those droplets are applicable for all impacting Newtonian fluids during inkjet printing:[7]

$$We = \frac{U^2 \rho L}{\gamma} \tag{S1}$$

When *We* << 1 (based on droplet radius), the spreading of droplets is capillarity-driven and the influence of initial droplet velocity is negligible. When *We* >> 1, the spreading of droplets is impact-driven and the influence of initial droplet velocity is significant. In addition, those authors characterized the resistance to spreading with *Oh* number (based on droplet radius). At very low values of *Oh* number, the spreading is resisted by inertial oscillations of droplets post impact. At very high values of *Oh* number, the spreading is resisted by viscosity of droplets.

Splashing of impacting droplets, which is the physical separation of a part of the fluid from droplets at the impact site,[8] decreases the success of inkjet printing. Mundo *et al*.[9] studied the onset of splashing of Newtonian fluids on a flat surface with the splash parameter ($K_{sp}$):

$$K_{sp} = (Oh^{-0.25})\ (We^{0.625}) \tag{S2}$$

The splash parameter *K* depends on surface roughness of substrates and for a flat-smooth surfaces $K_{sp}$ is 57.7 (based on droplet diameter). When *Ksp* > 57.7, impinging droplets splash on a smooth surface. When $K_{sp}$ < 57.7, impinging droplets do not splash and are completely deposited on a smooth surface. Although, elasticity influences the spreading behavior of droplets for non-

Newtonian viscoelastic fluids and suppresses splashing,[10] a detailed analysis of it is beyond the scope of this work.

An operating map which is presented in Figure S11 was prepared to predict the spreading behavior of impacting droplets of surfactant-free protein solutions on a dry solid surface. The map was constructed with the droplet velocity and the dimensionless numbers representing the fluid properties of protein solutions (see Table S2 for values of *Oh* and *We* numbers). The spreading of impacting droplets of collagen solutions with concentrations of 0.3 and 0.5 mg/mL is impact-driven and controlled by the droplet velocity according to the map. At the same time, the spreading of those droplets is resisted by the inertial oscillations of the droplets post impact. The impacting droplets of fibrinogen solutions with concentrations of 5, 10, and 20 mg/mL and thrombin solutions with concentrations of 5, 10, 15, 20 U/mL show similar spreading behavior according to the map. In addition, the droplets of those protein solutions do not splash upon impacting a dry solid surface. To confirm this, the droplets of 0.5 mg/mL collagen, 10 mg/mL fibrinogen, and 20 U/mL thrombin solutions were deposited on a dry glass slide by moving the inkjet nozzle with a velocity of 1 mm/s across the slide. The droplets were completely deposited on the slide and did not splash upon impact (see the inset in Figure S11). Hence, the droplets of dilute collagen, fibrinogen, and thrombin solutions do not splash upon impacting a dry solid surface and their spreading is controlled by their velocity.

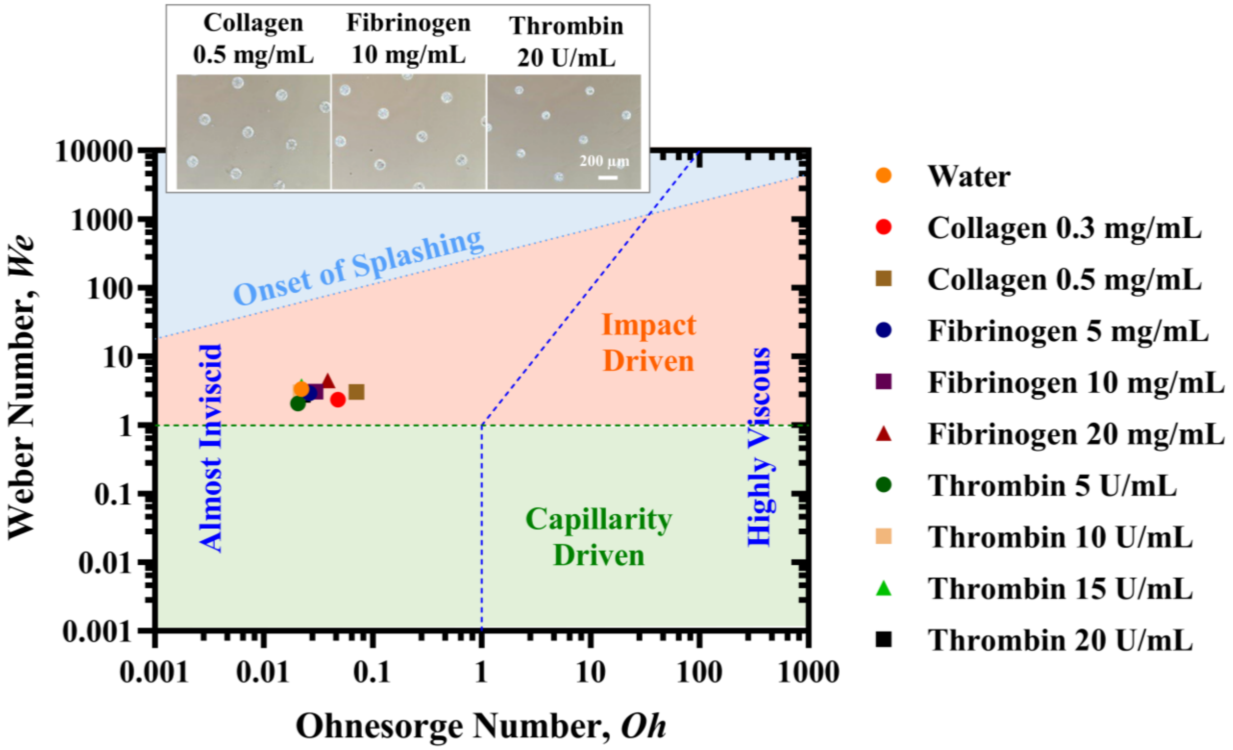

**Figure S11.** Map for predicting spreading behavior of impacting droplets of surfactant-free dilute collagen, fibrinogen, and thrombin solutions on a dry solid surface was constructed with the dimensionless numbers representing the fluid properties of those solutions, drawn following the schematic of Schiaffino *et al*.[6] The dimensionless numbers were calculated with the droplet radius and accordingly splash parameter $K_{sp}$, which was originally calculated with the droplet diameter in the literature, was adjusted. Inset shows the microscopic images of droplets of 0.5 mg/mL collagen, 10 mg/mL fibrinogen, and 20 U/mL thrombin solutions deposited on dry glass slides.

**Table S1.** Dimensionless parameters and time scales associated with breakup of droplets during DOD inkjet printing of surfactant-free protein solutions.

| Fluid | Density ($kg/m^3$) | Surface Tension (mN/m) | Viscosity (mPa.s) | Droplet Radius (μm) | Velocity (m/s) | *c/c** | *Oh* | *Z* | $t_c$ (μs) | $\tau_c$ (μs) | $\tau_v$ (μs) | $l_{sec}$ (nm) | $t_{sec}$ (ns) |
|---|---|---|---|---|---|---|---|---|---|---|---|---|---|
| Milli-Q Water | 1008 | 72 | 0.99 | 29.5 | 2.70 | 0.00 | 0.011 | 95 | 109 | 55 | 0.82 | 13 | 0.2 |
| Collagen 0.3 mg/mL | 965 | 68 | 2.25 | 33.5 | 2.19 | 0.60 | 0.025 | 39 | 131 | 55 | 1.98 | 77 | 3 |
| Collagen 0.5 mg/mL | 958 | 65 | 3.13 | 31.2 | 2.54 | 1.00 | 0.036 | 28 | 129 | 56 | 2.90 | 158 | 8 |
| Collagen 1 mg/mL | 927 | 64 | 5.74 | n/a | n/a | 1.99 | 0.068 | 15 | 140 | 56 | 5.39 | 556 | 50 |
| Collagen2 mg/mL | 923 | 58 | 16.41 | n/a | n/a | 3.99 | 0.205 | 5 | 131 | 59 | 16.96 | 5025 | 1420 |
| Fibrinogen 5 mg/mL | 936 | 51 | 0.96 | 28.0 | 2.34 | 0.13 | 0.013 | 79 | 109 | 63 | 1.13 | 19 | 0.4 |
| Fibrinogen 10 mg/mL | 921 | 47 | 1.05 | 28.9 | 2.29 | 0.27 | 0.015 | 68 | 110 | 65 | 1.36 | 26 | 0.6 |
| Fibrinogen 20 mg/mL | 929 | 45 | 1.31 | 27.3 | 2.78 | 0.53 | 0.019 | 54 | 109 | 67 | 1.75 | 41 | 1 |
| Fibrinogen 50 mg/mL | 978 | 45 | 2.48 | n/a | n/a | 1.34 | 0.034 | 29 | 145 | 69 | 3.31 | 140 | 8 |
| Thrombin 5 U/mL | 989 | 69 | 0.89 | 27.5 | 2.25 | $3\times10^{-4}$ | 0.010 | 101 | 110 | 56 | 0.78 | 12 | 0.2 |
| Thrombin 10 U/mL | 988 | 59 | 0.89 | 28.6 | 2.46 | $6\times10^{-4}$ | 0.011 | 94 | 110 | 60 | 0.91 | 14 | 0.2 |
| Thrombin 15 U/mL | 983 | 58 | 0.89 | 24.2 | 2.95 | $9\times10^{-4}$ | 0.011 | 93 | 106 | 60 | 0.92 | 14 | 0.2 |
| Thrombin 20 U/mL | 968 | 58 | 0.90 | 27.3 | 2.43 | $12\times10^{-4}$ | 0.011 | 92 | 108 | 60 | 0.92 | 14 | 0.2 |

*c/c** – Overlap concentration of proteins in Dulbecco's phosphate buffered saline solution.
*Oh* – Ohnesorge number based on nozzle diameter.
*Z* – Inverse Ohnesorge number based on nozzle diameter.
$\tau_c$ – Inertio-capillary time scale based on nozzle diameter.
$\tau_v$ – Visco-capillary time scale based on nozzle diameter.
$l_{sec}$ – Viscous length scale of secondary microthread based on nozzle diameter.
$t_{sec}$ – Viscous time scale of secondary microthread based on nozzle diameter.

**Table S2.** Dimensionless parameters associated with impinging droplets during DOD inkjet printing of surfactant-free protein solutions.

| Fluid | Density (kg/m$^3$) | Surface Tension (mN/m) | Viscosity (mPa.s) | Droplet Radius (µm) | Velocity (m/s) | $Oh_{Drop}$ | $We_{Drop}$ |
|---|---|---|---|---|---|---|---|
| Milli-Q Water | 1008 | 72 | 0.99 | 29.5 | 2.70 | 0.021 | 2.99 |
| Collagen 0.3 mg/mL | 965 | 68 | 2.25 | 33.5 | 2.19 | 0.048 | 2.28 |
| Collagen 0.5 mg/mL | 958 | 65 | 3.13 | 31.2 | 2.54 | 0.071 | 2.96 |
| Collagen 1 mg/mL | 927 | 64 | 5.74 | n/a | n/a | n/a | n/a |
| Collagen2 mg/mL | 923 | 58 | 16.41 | n/a | n/a | n/a | n/a |
| Fibrinogen 5 mg/mL | 936 | 51 | 0.96 | 28.0 | 2.34 | 0.026 | 2.84 |
| Fibrinogen 10 mg/mL | 921 | 47 | 1.05 | 28.9 | 2.29 | 0.030 | 2.99 |
| Fibrinogen 20 mg/mL | 929 | 45 | 1.31 | 27.3 | 2.78 | 0.039 | 4.35 |
| Fibrinogen 50 mg/mL | 978 | 45 | 2.48 | n/a | n/a | n/a | n/a |
| Thrombin 5 U/mL | 989 | 69 | 0.89 | 27.5 | 2.25 | 0.021 | 2.00 |
| Thrombin 10 U/mL | 988 | 59 | 0.89 | 28.6 | 2.46 | 0.022 | 2.93 |
| Thrombin 15 U/mL | 983 | 58 | 0.89 | 24.2 | 2.95 | 0.024 | 3.57 |
| Thrombin 20 U/mL | 968 | 58 | 0.90 | 27.3 | 2.43 | 0.023 | 2.68 |

$Oh_{Drop}$ – Ohnesorge number based on drop radius.
$We_{Drop}$ – We number based on drop radius.
